# Distinct neural processes link speech planning and execution

**DOI:** 10.1101/2024.10.07.617122

**Authors:** Suseendrakumar Duraivel, Shervin Rahimpour, Katrina Barth, Chia-Han Chiang, Charles Wang, Stephen C. Harward, Shivanand P. Lad, Daniel P. Sexton, Allan H. Friedman, Saurabh R. Sinha, Gregory Hickok, Derek G. Southwell, Jonathan Viventi, Gregory Cogan

**Affiliations:** Department of Biomedical Engineering, Duke University, Durham, NC, USA; Department of Neurosurgery, Duke School of Medicine, Durham, NC, USA; Duke Comprehensive Epilepsy Center, Duke School of Medicine, Durham, NC, USA; Penn Epilepsy Center, Perelman School of Medicine, University of Pennsylvania, Philadelphia, PA, USA; Department of Cognitive Sciences, University of California, Irvine, CA, USA; Department of Language Science, University of California, Irvine, CA, USA; Department of Neurobiology, Duke School of Medicine, Durham, NC, USA; Department of Neurology, Duke School of Medicine, Durham, NC, USA; Department of Psychology and Neuroscience, Duke University, Durham, NC, USA; Center for Cognitive Neuroscience, Duke University, Durham, NC, USA

**Keywords:** Speech, IEEG, syllable sequences, speech planning, speech production

## Abstract

Speaking is the primary way that humans communicate. This communication is enabled by a production system that can plan and execute unique combinations of speech sounds. Although a distributed network of brain regions has been implicated in speaking, it is unclear how planning and execution of speech are coordinated to produce meaningful sounds. Leveraging the high spatio-temporal resolution of intracranial recordings at different spatial scales, we show distinct neural mechanisms that facilitate speech planning and execution. During planning, different levels of speech units are coded discretely at distinct prefrontal sites. These planned units are then dynamically integrated at various cortical levels to guide subsequent execution. During speech execution, speech motor regions generate continuous sequences that reflect both discrete speech sound units and their transitional properties between units. This rapid neural transition from discrete speech units to motor sequences links speech planning with execution and enables our effortless ability to speak.

## Introduction

Speech production enables verbal communication through planning and motor execution of combinations of speech sounds^1–3^. This fundamental human capacity of speech occurs by turning mental ideas to words^4–6^, which are then mapped to smaller speech units to generate motor sequences for execution^7,8^. This ability is acquired over years of learning to speak in which speech plans organize the vocal execution for continuous speech production. Despite this effortless everyday usage, the neural basis of how speech plans and execution are organized and linked is not well characterized.

From the psycholinguistic literature^9–11^, speech plans are hypothesized to operate at two distinct levels: a framing level which corresponds to the syllable, and a content level which combines phonemes within syllable frames^12–16^. Behavioral evidence for these planning elements comes from analyses of speech errors such as slips of the tongue and phoneme exchange errors during everyday speech and in aphasia, suggesting neural processes for distinct stages in speech planning^9,17,18^. These error patterns reveal that different linguistic units – phonemes and syllables, – interact systematically during speech planning, suggesting neural substrates for distinct stages in this pre-execution process.

Planned utterances are subsequently fed into a motor execution system that: 1) generates commands to control the sequence of vocal tract gestures for articulation, and 2) updates the commands based on sensory feedback from the auditory and the somatosensory system^12–15,19,20^. Together, representations for syllables and phonemes combine during planning and execution to allow the speaker to effortlessly utter any sequence of speech sounds within the phonological rules of their proficient language^9–11^. Despite the importance of planning and execution for speech production, the neural processes that distinguish these hypothesized distinct stages remain largely unexplored (**Fig. 1a**).

**Figure 1.**
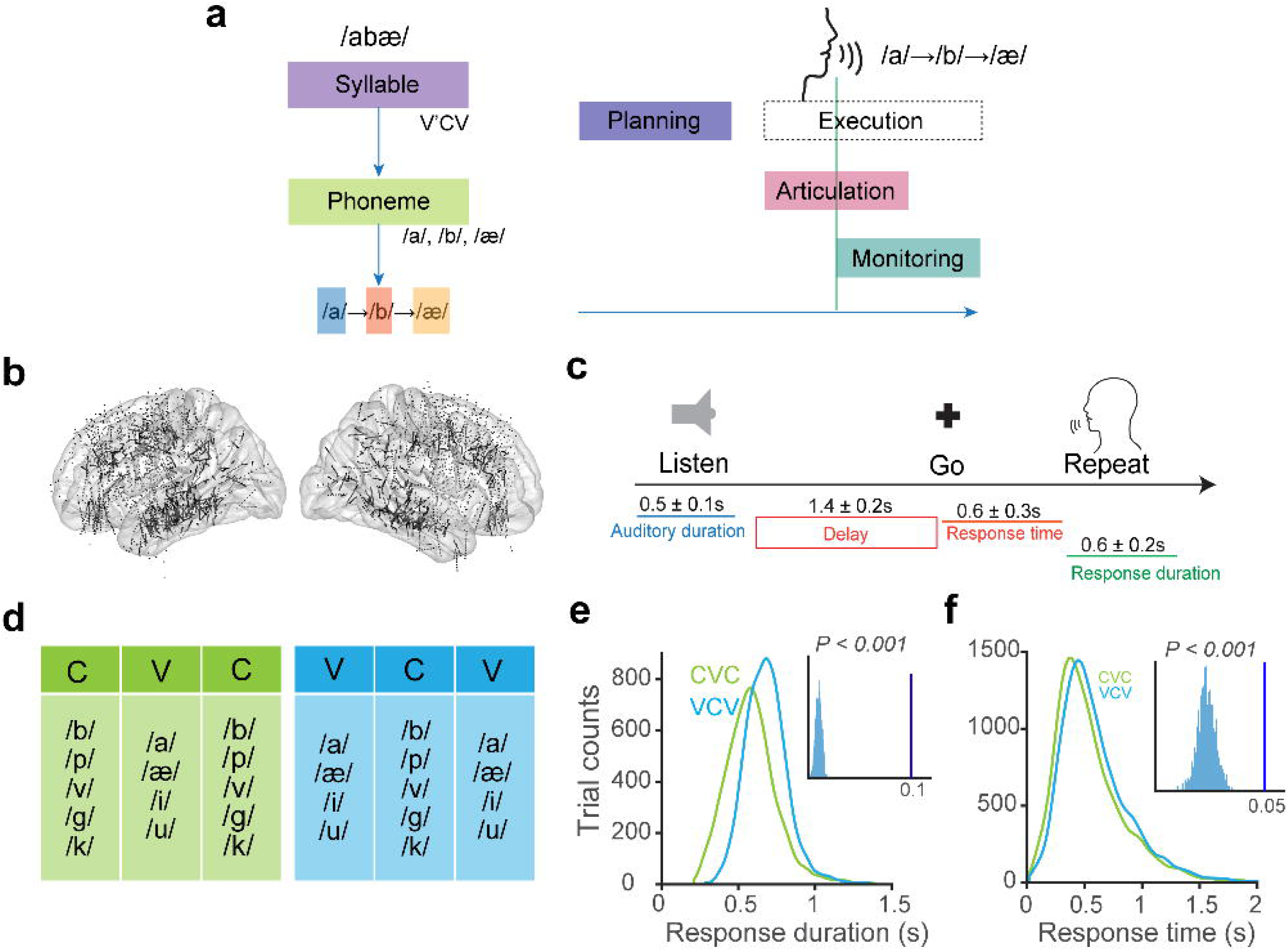
Pseudo-word repetition task to study planning and execution for speech production. **a.** Speech planning and execution at the sublexical level (post word formation) involves the composition of a syllable frame and the hierarchical incorporation of phonemes within this frame for the execution of the ordered speech sequence. Graphical representation of speech neural mechanisms from the planning of intended speech units to the execution of speech sound sequences. Speech production involves planning prior to execution, and execution which involves motor articulation followed by auditory monitoring of vocal feedback. **b.** Location of all 8106 recorded intracranial electrodes from 52 patients in both left and right hemispheres. **c.** Schematic of pseudo-word repetition task. Color bars indicate the duration of the auditory stimulus (blue), response time from the Go cue (orange), and response duration (green). **d.** Pseudo-word tokens used for the speech repetition task differed on their syllabic frame: consonant-vowel-consonant (CVC, green) and vowel-consonant-vowel (VCV, blue). The phonemes were chosen from a fixed for distribution for both consonants (5) and vowels (9) at each position within the pseudo-word. **e.** Histogram distributions of response duration for both CVC and VCV trials. The durations of VCV trials were significantly longer than the CVC trials (linear mixed effects [LME] modeling, t_(8696)_ = 10.7 β = 0.098, *p < 0.001*, 95% CI = 0.08 to 0.1, two-sided). The dark blue line indicates the beta estimate of the actual LME model, and the histograms (light blue) indicate the estimated distributions from permuted models (1000 permutations). **f.** Histogram distributions of response times for both CVC and VCV trials. The response time of CVC trials were significantly faster than the VCV trials (linear mixed effects [LME] modeling, t_(8698)_ = 3.7, β = 0.045, *p < 0.001*, 95% CI = 0.02 to 0.07, two-sided). The dark blue line indicates the estimate of the actual LME model, and the histograms (light blue) indicate the estimate distributions from permuted models (1000 permutations). Longer response times for di- vs. mono- syllabic utterances suggest that the syllable frame is used as a sub-lexical unit for speech planning.

Previous work attempting to distinguish planning and execution of speech production have come from neuroimaging studies, which have identified distributed neuroanatomical networks of frontal, parietal, and temporal regions. Specifically, these regions include: inferior frontal gyrus (IFG), rostral and caudal middle frontal gyrus (rMFG & cMFG), precentral and postcentral gyrus (PrCG & PoCG; sensorimotor cortex), supramarginal gyrus (SMG), inferior parietal cortex (IPC), insula, superior temporal gyrus (STG), and primary auditory cortex (PAC)^1,12,21–30^. Within this network, prefrontal regions have primarily been associated with planning, sensorimotor circuits (PrCG, PoCG, SMG, and IPC) associated with motor execution,^3,31^ and superior temporal regions and the insula associated with monitoring based on sensory feedback (auditory and somatosensory)^3,32–34^, as well as internal error correction^35,36^. Significant progress has been made to understand how speech is executed by sensorimotor circuits through coordinated activity to produce speech sounds^4,24,29^, as well as how speech is monitored through neural suppression of self-generated speech sounds in STG^37–40^. For speech planning, however, neuroimaging studies have been limited by their lack of precise timing to distinguish neural processes associated with planning from neural processes associated with execution. This limitation makes it difficult to directly study the presence of distinct speech units such as syllables and phonemes, as distinct processing stages during planning.

One indirect approach to study the structure of speech planning has been to manipulate aspects of syllables (order, complexity, and novelty of syllables), and examine the cortical regions activated using neuroimaging^41^. Notably, studies that have manipulated syllable structures and found activation in the posterior inferior frontal gyrus (pars-opercularis)^10,11,41–43^, supporting the idea that the left IFG could be involved in syllable planning for speech production. However, despite these insights, the precise temporal dynamics involved in planning and execution across the speech network, particularly how syllable and phoneme processes unfold over time – remains unknown.

Our understanding of the temporal dynamics of speech planning and execution have mostly come indirectly from computational models of speech production, including the DIVA (Directions Into Velocities of Articulators), GODIVA (Gradient Order DIVA), and hierarchical state feedback control (HSFC) models have been developed based on the neuroimaging literature to integrate psycholinguistic (syllable-phoneme) and motor-control (planning-execution) theories that model the accurate planning, execution, and monitoring of sequences of speech sounds^14,16,27^. The DIVA model provides a neuroanatomically explicit framework for speech motor control through parallel feedforward and feedback systems, mapping abstract speech representations to articulatory movements with error correction via auditory and somatosensory feedback^27^. GODIVA extends DIVA by adding mechanisms for sequencing syllables and phonemes in multi-syllabic utterances by incorporating sequential structure buffers and phonological content buffers^16^. Further, Hickok’s HSFC model integrates these approaches within an hierarchical architecture where higher-level control codes speech segments at syllabic level while the lower-level execution processes update the contents at phonemic level^14^. Simulations using this hierarchical organization resulted in accurate generation of phoneme sequences, with a higher order planning state coding speech at the syllabic level and a lower order planning state at the phoneme level.^13,14^ These computational models suggest that during planning, two distinct speech representations (syllables and phonemes) are composed both hierarchically and in serial order, with syllables planned prior to phoneme plans. While these models describe the computational and anatomical structure of speech production and incorporate both feedforward and feedback control mechanisms, they cannot empirically specify the precise timing and temporal evolution of neural activations across regions during natural speech production. As a result, direct empirical evidence for the temporal ordering and integration of syllable and phoneme representations during planning and execution is lacking, due to limitations in recording technology for resolving neural signals with both high spatial and temporal precision.

Intracranial recordings using electrocorticography (ECoG) and stereo-electroencephalography (SEEG) record neural activity directly from cortex with high temporal resolution (milliseconds) with precise anatomical localization^44–47^. This technology has been further advanced through the development of high-density micro-electrocorticographic (µECoG; 1.33 – 1.72 mm inter-electrode distance) arrays that can measure micro-scale cortical activity with millimeter scale spatial resolution^48–52^. Neural activations from ECoG and SEEG in the high-gamma band (HG: 70-150 Hz) have been shown to correlate with multi-unit firing rates, making this signal a good measure of local neural activity^53–57^.

During speech production, studies have shown HG activations in the ventral sensorimotor cortex which encode articulatory properties during speech motor execution^19,20,58–63^. These findings have provided detailed insights into how specific brain regions control vocal tract movements and coordinate articulatory gestures. However, fundamental questions remain about how the brain enables the transition from speech planning to precise motor execution.

Surprisingly, there has been limited intracranial work examining speech planning, and the critical transition between planning and execution. One previous study used overt sentence completion to examine lexical access in the IFG prior to articulation^64^, but this activity could be related to either speech perception or production processes, making its role in speech production unclear. Another study used an unstructured experimental setting to examine the timing of planning in relation to turn taking in conversation^65^ but it did not examine the content of planning or provide evidence for how speech plans enable motor execution. A more recent study demonstrated that speech motor sequencing relies on a distributed network centered on the middle precentral gyrus, showing sustained activity that is modulated by sequence complexity, yet little is discussed about the presence or absence of hierarchical organization of sublexical units (syllables and phonemes), or the temporal dynamics of the integration during the planning-to-execution transition^66^. These studies do highlight the promise of intracranial recordings for studying different stages of speech production via the measurement of spatially specific neural representations. The high temporal and spatial resolution of intracranial recordings also presents an opportunity to delineate the neural substrates associated with the transition from speech planning to execution. Understanding this transition could have important implications for treating neurological conditions that affect both higher order cognition and speech production, as well as in developing more advanced neural prosthetic technology.

To elucidate the neural substrates for planning and execution in speech production, we performed intracranial recordings with 52 patients while they verbally articulated constructed pseudo-words. We show distinct neural activations for planning and execution that represent both syllables and phonemes. The neural activations demonstrate distinct spatial and temporal patterns that link speech planning to execution. During the transition from planning to execution, we showcase neural processes that transform speech sound units - syllables and phonemes - from plans to motor execution. In execution, we show evidence for motor sequencing through ordered patterns of phonemes as well as evidence for the neural processing of the transitions between phonemes. Together, these findings provide evidence for distinct neural patterns that enable the rapid transition from discrete speech units during planning to sequenced speech sounds during motor execution, supporting fluent speech production.

## 1 Results

We performed a speech task with 52 patients (n = 52, mean age = 33.1 years, 23 females, Supplementary table 1) in the epilepsy clinical monitoring unit, and with 3 patients intra-operatively (n = 3, mean age = 64.3 years, 0 females, Supplementary table 1). Patients in the monitoring unit were implanted with clinical intracranial electrodes (electrocorticography [ECoG: 2 patients] or stereo-electroencephalography [SEEG: 50 patients]) for the localization of epileptic tissue to guide surgical resection or neuromodulation targeting (**Fig. 1b**). Patients recorded intra-operatively were awake as part of their surgical procedure for tumor resection (1 patient) or deep brain stimulator (DBS) placement for the treatment of movement disorders (2 patients), and these neural recordings were obtained at the micro-scale cortical level with high spatial density using µECoG electrode arrays^63^.

Participants performed a controlled pseudo-word repetition task. Pseudo-words were presented auditorily through speakers connected to a presentation laptop running Psychtoolbox^67^. After the pseudo-word was presented, there was a short delay (1.2 – 1.6 s), followed by a prompt (using a visual “Speak” cue), during which the patients were instructed to repeat back the presented pseudo-word stimulus (**Fig. 1c**). Each pseudo-word was either composed with a monosyllabic (consonant-vowel-consonant: CVC) or a disyllabic (vowel-consonant-vowel: VCV) structure, that contained three phonemes (out of a set of 9 total available phonemes - **Fig. 1d**, Supplementary table 2). Each patient completed four task blocks that contained 52 unique tokens each. Due to the shorter availability of time, patients in the intra-operative setting underwent a similar but modified version of the pseudo-word repetition task in which they repeated the pseudo-words immediately after the auditory presentation. The intraoperative patients completed three to four repetitions of task blocks.

### 1.1 Behavioral evidence for syllables

On average, the patients who underwent delayed speech-repetition task, repeated the pseudo-words after 0.58 seconds (standard deviation: ± 0.33 s) from the visual “Speak” cue and with an average duration of 0.65 seconds (± 0.16 s). Mean response times (RT) and response durations (RD) were (mean ± SD): CVC: RT = 0.56 ± 0.35 s, RD = 0.58 ± 0.17 s; and VCV: RT = 0.6 ± 0.32 s, RD = 0.68 ± 0.14 s (**Fig. 1e, f**).

To quantify the effect of differences in syllable on behavioral performance, we performed linear mixed effects (LME) modeling on each trial, with a fixed effect modeling the syllable identity linked to each pseudo-word and a random effect modelling the patient identity. VCV structure utterances had longer durations compared to CVC structures [LME: t_(8696)_ = 10.7, β = 0.098, *p < 0.001*, 95% CI = 0.08 to 0.1, **Fig. 1e**, Supplementary Fig. 1], in keeping with their longer articulatory and acoustic duration (2 vs. 1 syllable). Interestingly, response times were significantly delayed for disyllabic VCV structures as compared to monosyllabic CVC structures [LME: t_(8698)_ = 3.7, β = 0.045, *p < 0.001*, 95% CI = 0.02 to 0.07, **Fig. 1e**, Supplementary Fig. 1], suggesting that speech motor planning reflects the composition of syllables. Further, this behavioral result also held when controlling for the response duration, supporting the interpretation that these effects reflect planning of syllables rather than the overall utterance length [LME: t_(9084)_ = 2.75, β = 0.103, *p = 0.006*, 95% CI = 0.03 to 0.18, Supplementary Fig. 1].

### 1.2 Neural processes for speech planning and execution

To investigate the neural processes involved in speech production, we first extracted neural activations in the high-gamma band (HG: 70 – 150 Hz) that have previously been shown to be an index of local neural computation and a correlate of multi-unit firing^54,56,57^. Only correctly articulated trials with response times greater than 50 ms (from “Speak”/go cue) were used for neural analysis. Out of 8106 preprocessed IEEG electrodes from 52 patients in the clinical monitoring unit, we observed significant increases in HG power in 3534 electrodes (2895 electrodes with grey matter contacts) during speech production as compared to a pre-stimulus baseline (one-sided permutation test with post-hoc FDR correction per patient, *p < 0.05*, **Fig. 2a**, Supplementary table 2). Speech neural activations were present in both the left (1749 electrodes) and right (1785 electrodes) hemisphere (**Fig. 2b**), confirming the bilateral nature of speech production networks^20^.

**Figure 2.**
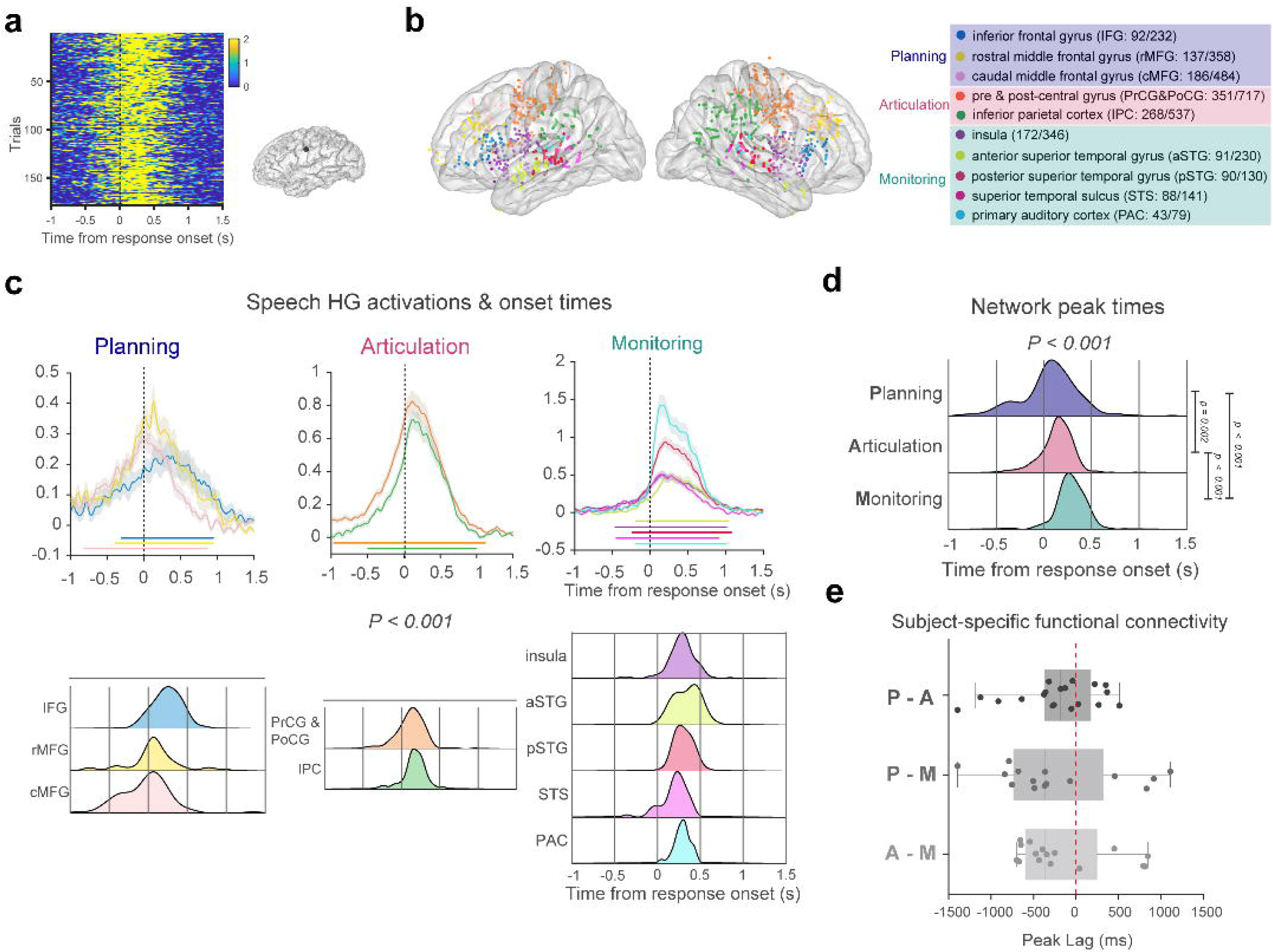
Cortical activations from intracranial recordings during speech production. **a.** Speech neural activations in the high-gamma band (HG: 70 – 150 Hz; z-score normalized with respect to baseline) of an example electrode in the pre-central gyrus that showed significant modulation during the speech utterance at the single trial level (FDR-corrected non-parametric permutation test). **b.** Location of all 3534 intracranial electrodes in both left (1749) and right (1785) hemisphere, with significant HG modulation during the speech utterance. The electrodes are color-coded with respect to anatomical regions-of-interest (ROIs), and are assigned to planning, articulation, and monitoring networks based on previous literature. The numbers in the brackets indicate the number of significant electrodes with respect to the total number of recorded electrodes per ROI. **c. Top:** Mean HG activations of ROIs during response onset in left hemisphere, averaged within patient. Shaded area denotes standard error. Colored horizontal bars indicate time-periods of significant activation above baseline (one-sided permutation test with cluster correction, cluster forming threshold *p<0.05,* 1000 permutations). **Bottom:** Electrode distributions of peak HG time-points per ROI in left hemisphere. Significant differences in peak activations were observed and there was a general trend of temporal progression of neural activations across ROIs around the speech utterance (one-way ANOVAs: F_9,717_ = 10.87, *p < 0.001,* two-sided). **d.** Electrode distributions of peak HG time-points per ROI groups. This temporal progression was also observed via significant differences in peak activations (one-way ANOVAs: F_2,724_ = 30.7, *p < 0.001,* two-sided). Post-hoc one-sided t-tests indicate pairwise significance between the speech networks in left hemisphere: T_Planning_ < T_Articulation,_ t_(365)_ = 2.938*, p* = 0.002; T_Articulation_ < T_Monitoring,_ t_(507)_ = 9.25*, p < 0.001*, T_Planning_ < T_Monitoring,_ t_(394)_ = 9.84*, p < 0.001*). **e**. Functional connectivity analysis validates the temporal progression of speech networks: peak lag of the net cross-correlation of the HG signals was computed within each subject. Dots represent individual subject median peak lags; box plots show the subject distribution of lags for each network pair comparison. The box spans the interquartile range (IQR), and whiskers extend to the observed value withing 1.5 × IQR of each quartile boundary. Negative lags indicate that the first network (e.g. planning) leads the second (e.g. articulation). In all three pairwise comparisons, real peak lags were significantly earlier than the control obtained under a circular-shift null (one-sided paired Wilcox signed-rank test): Planning-Articulation (n=19 subjects, median = -180ms, W = 36, *p = 0.007*); Articulation-Monitoring (n=16, median = -398 ms, W = 16, *p = 0.003*); and Planning-Monitoring (n=15, median = -370ms, W = 30, *p = 0.047*). Together, these analyses demonstrate the temporal progression of neural responses for planning, articulation, and monitoring for speech production in the left hemispheric speech network.

We further selected electrodes within anatomical regions of interest (ROIs) that have been shown to be active for speech production^4,16,20,65,68^. The selected ROIs in both hemispheres included: inferior frontal gyrus (IFG: 92 significant electrodes), rostral middle frontal gyrus (rMFG: 137), caudal middle frontal gyrus (cMFG: 186), precentral and postcentral gyrus (PrCG & PoCG: 351), inferior parietal cortex (IPC: 268), insula (172), anterior superior temporal gyrus (aSTG: 91), posterior superior temporal gyrus (pSTG: 90), superior temporal sulcus (STS: 88), and primary auditory cortex (PAC: 43). See supplementary table 3 for the anatomical assignment.

To characterize the timing of activation between ROIs, we estimated the peak onsets of HG activation for each electrode relative to the utterance onset (**Fig. 2c**). There were significant differences in peak activations between ROIs in both the left and right hemispheres (one-way ANOVA; left hemisphere: F_9,717_ = 10.87, *p < 0.001*, right hemisphere: F_9,781_ = 6.69, *p < 0.001.* **Fig. 2c**, Extended data fig. 1). Based on neuro-anatomical studies that investigated speech articulation^4,16,20,65,68^, we divided these ROIs into three distinct networks based on previous literature: planning (IFG, rMFG, and cMFG), articulation (PrCG & PoCG, and IPC), and monitoring (aSTG, pSTG, STS, and PAC). As hypothesized, we found that in the left hemisphere, electrodes in the planning network were activated early, followed by electrodes in the articulation network, and then the monitoring network (one-way ANOVA: F_2,724_ = 30.7, *p < 0.001,* **Fig. 2d**). Pair-wise post-hoc t-tests showed that the mean values were significantly different between all three groups, supporting distinct functional roles rather than simple anatomical groupings (*p < 0.05,* T_Planning_ < T_Articulation,_ t_(365)_ = 2.938, p = 0.002; T_Articulation_ < T_Monitoring,_ t_(507)_ = 9.25, *p < 0.001*, T_Planning_ < T_Monitoring,_ t_(394)_ = 9.84, *p < 0.001*, **Fig. 2d**). Differences were also significant in the right hemisphere (F_2,788_ = 8.86, *p < 0.001,* T_Articulation_ < T_Monitoring,_ t_(528)_ = 7.53 *p < 0.001*, T_Planning_ < T_Monitoring,_ t_(385)_ = 2.36*, p* = 0.019, Extended data fig. 1). However, the prefrontal-planning activations occurred later in time than the articulation regions (T_Planning_ > T_Articulation,_ t_(511)_ = 2.3*, p* = 0.021), which is consistent with the role of the right hemisphere in error monitoring^69,70^. A similar organization existed when the neural data were aligned to the go-cue (Supplementary Fig. 2).

We next sought to investigate whether these speech networks were functionally connected during speech production. We computed the cross-correlation between network-averaged HG time-courses between all pairs of speech networks, within each subject (**Fig. 2e**). In all three pairwise comparisons, the cross-correlation peak lags were significantly more negative (against circular-shift null distribution) when the networks were arranged with respect to the purported temporal order (one-sided Wilcox signed-rank test): planning preceded articulation by a median of 180 ms (n = 19 subjects, W = 36*, p = 0.007*); articulation preceded monitoring by a median of 398 ms (n = 16 subjects, W = 16*, p = 0.003*); and planning preceded monitoring by a median of 370 ms (n = 15 subjects, W = 30*, p = 0.047*). This presence of functional connectivity indicates that the network activations provide evidence for a temporal order of speech information within-subject.

While these ROI-based network assignments were grounded in established neuroanatomical literature, such a-priori groupings could potentially introduce bias into the analysis. To address this potential limitation and validate the network organization in a data-driven manner, we performed non-negative matrix factorization (NNMF) on the z-scored HG activation matrix to identify latent neural speech components without prior assumptions (Supplementary Fig. 3). The resulting five temporal components revealed distinct activation profiles that closely matched our hypothesized speech production stages (Supplementary Fig. 3b): Cluster 1 and 4 captured early and late auditory monitoring respectively, Cluster 2 showed articulation-related activity, Cluster 3 reflected planning processes, and Cluster 5 indicated sustained verbal working memory maintenance, as part of the sensory-motor transformation. Critically, on examining how these data-driven clusters mapped onto the anatomically defined ROIs and speech networks, we found strong correspondence between the unsupervised decomposition and our ROI-based groupings. Planning regions (IFG, rMFG, cMFG) showed maximal loading on Cluster 3, articulation regions (PrCG, PoCG, IPC) aligned predominantly with Cluster 2, and monitoring regions (aSTG, pSTG, STS, & PAC) correspond to Clusters 1 and 4. This convergence validates the network organization and demonstrates that the temporal progression reflects functional specialization rather than arbitrary anatomical clustering. Overall, these results clearly demonstrate the presence of three distinct speech networks in the left hemisphere: 1) a planning network in the pre-frontal regions to initiate speech, 2) an articulation network that generates motor commands and executes planned speech, and 3) a monitoring network for auditory feedback tracking.

### 1.3 Neural syllabic code is present prior to articulation

We next sought to investigate the neural basis for speech representation associated with the three levels of processing (planning, articulation, and monitoring) at each stage. Based on prior computational work, we theorized that speech production involves the recruitment of syllables during the planning stage^4,11,14,42^, followed by their execution during the produced utterance (articulation and monitoring)^14,27^. To assess the specific timing of neural activations that reflect differences in the syllable structure of the utterances, we examined differences in ROI-grouped HG activations between syllables (ΔHG; VCV and CVC pseudo-word trials, see Methods), at every time point around the response epoch (**Fig. 3a**). With phonemes controlled at each position within the syllable, these HG contrasts (ΔHG) can reveal activation patterns selective to the syllable alone. Results show that neural activations associated with the syllable structure showed temporal components with significant syllable coding (time clusters of significant differences in activation between VCV and CVC) for each speech network in the left hemisphere. We observed the earliest coding (ΔHG) in the prefrontal-planning network (cluster onset: -0.73s, peak onset: -0.16s), followed by the articulatory network (cluster onset: -0.22s, peak onset: 0.23s), and monitoring network (cluster onset: -0.16s, peak onset: 0.32s (one-way ANOVA, F_2,495_ = 64.3, *p < 0.001*, **Fig. 3b**). Post-hoc t-tests showed that the differences showed pairwise significance between the three areas (p < 0.05), except for between the articulatory and the monitoring areas, confirming the temporal progression from syllable planning to their motor execution (T_Planning_ < T_Articulation,_ t_(333)_ = 3.3*, p = 0.001*; T_Articulation_ < T_Monitoring,_ t_(403)_ = 1.3*, p = 0.2* (not significant), T_Planning_ < T_Monitoring,_ t_(352)_ = 3.78*, p < 0.001*, **Fig. 3c**). To directly investigate this ROI division of syllable coding, we repeated the analysis on electrodes grouped within each ROI (Extended data fig. 2a, except STS; p < 0.05, one-sided permutation test with cluster correction). The earliest time clusters for syllable contrasts were observed in left hemispheric planning regions (cluster start time & mean peak onsets): 1) IFG: -0.74s & -0.3s, 2) rMFG: -0.68s & -0.05s, and 3) cMFG: -0.7s & -0.17s. Speech articulatory ROIs demonstrated relatively late timing: 1) PrCG & PoCG: -0.6 & 0.83s, and 2) IPC: -0.1s & 0.3s. Finally, the monitoring ROIs exhibited timing closer to voice-onset times: 1) Insula: -0.27s & 0.32s, 2) aSTG: -0.22s & 0.37s, 3) pSTG: -0.16s & 0.26s, and 4) PAC: -0.16s & 0.31s. We further performed a one-way ANOVA to assess the temporal properties of syllabic coding, which demonstrated significant differences in peak coding between individual ROIs (Extended data fig. 2b, except STS, F_8,489_ = 18.1, *p < 0.001*)

**Figure 3.**
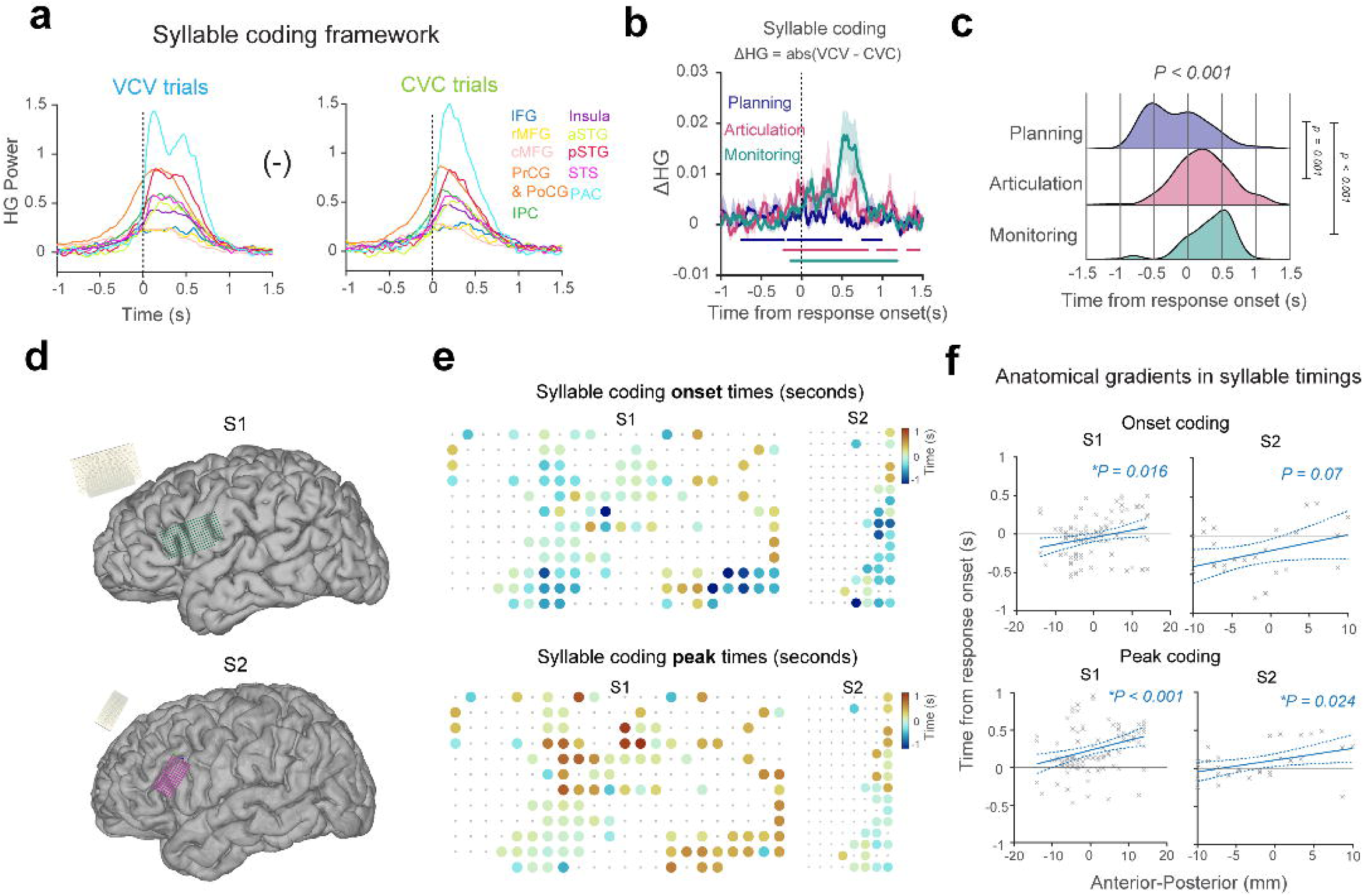
Neural code for syllables in the left hemisphere. Syllables are theorized to be composed during planning and used as frames for articulation and monitoring networks during speech execution**. a**. We analyze the neural code for the syllable frame using a syllable discrimination framework: ROI-grouped HG activations of VCV trials were contrasted (subtracted) with CVC trials. **b.** This analysis was carried out for each speech network and time points with significant syllable discrimination were found for planning, articulation and monitoring networks. Plots show mean ΔHG contrasts of VCV versus CVC for each network aligned to the response onset. Shaded area denotes standard error. Horizontal bars indicate durations of significant contrasts (across vs. within frame, one-sided permutation test with cluster correction, cluster forming threshold *p<0.05,* 1000 permutations). **c.** Histograms show electrode distributions of peak ΔHG contrasts per network: planning, articulation, and monitoring. Significant differences in peak ΔHG (one-way ANOVA, F_2,495_ = 64.3, *p < 0.001,* two-sided), followed by post-hoc one-sided t-tests indicate pairwise significance in syllabic frame contrasts (ΔHG) between the networks indicate a temporal progression of syllable activations across ROI networks (T_Planning_ < T_Articulation,_ t_(333)_ = 3.3*, p* = 0.001; T_Articulation_ < T_Monitoring,_ t_(403)_ = 1.3*, p* = 0.2 (not significant), T_Planning_ < T_Monitoring,_ t_(352)_ = 3.78*, p < 0.001*). **d.** µECoG captures spatio-temporally selective syllabic coding sites for speech planning and execution. High-density µECoG electrode arrays were implanted over planning (anterior) and execution (posterior) networks in two awake patients. Electrode arrays had either 256 electrodes with 1.72 mm inter-electrode distance for S1 (green) or 128 electrodes with 1.33 mm inter-electrode distance for S2 (pink). **e.** Neural code for the syllable was extracted by contrasting HG activations per syllabic structure (ΔHG: VCV – CVC) for each electrode. Heat maps show spatial activation patterns of onset and peak times of ΔHG activations for electrodes with significant syllable contrasts. Large circles reflect electrodes with significant syllable contrasts (one-side permutation test with cluster correction, cluster forming threshold *p < 0.05,* 1000 permutations, S1: 100/201, S2: 37/63). Non-significant electrodes are indicated by grey dots. These significant contrasts indicate syllable coding on an electrode specific level and demonstrate spatially selectivity on the underlying neuroanatomy. **f.** A spatial correlation analysis of the syllable contrast onsets along the anterior-posterior axis shows a spatial progression of temporal coding for syllable codes starting in the anterior planning section and moving towards the posterior execution network (S1: R^2^ = 0.05, F_2,92_ = 6.02, *p = 0.016*, S2: R^2^ = 0.08, F_2,28_ = 6.02, *p = 0.07,* two-sided). This spatio-temporal progression was also seen in the peak times (F-statistic test against the constant model, S1: R^2^ = 0.09, F_2,98_ = 10.9, *p < 0.001*, S2: R^2^ = 0.13, F_2,28_ = 5.68, *p = 0.024,* two-sided). Dotted lines indicate the 95% confidence interval of the linear model fit. The presence of a spatial gradience, extracted using high-density µECoG electrode arrays, demonstrate a potential anatomical substrate for the transition from planning to execution for speech production.

In the right hemisphere, significant differences in syllable neural codes were observed between the articulation and monitoring network (negative t-statistic on comparisons against planning network, T_Planning_ < T_Articulation,_ t_(406)_ = -5.4*, p < 0.001*, Supplementary Fig. 4), further, supporting the right hemisphere’s involvement in execution. Similar temporal progression is also observed when neural data was aligned with the go-cue onset, showing that the transition from planning to execution of syllables is also preserved at the commencement of planning with respect to the instructional cue (Supplementary Fig. 5). To address a potential confound between syllable class (mono-syllabic/CVC vs. di-syllabic/VCV) and initial phoneme category (C vs. V), we performed a control analysis by comparing the production of CV’CVC and CVC structures that differ in syllable number while sharing the same initial consonantal phoneme structure. We find that planning of monosyllabic vs. disyllabic utterances show similar planning and execution temporal activation patterns as in our main analysis (Supplementary Fig. 6). This analysis ruled out initial phoneme category influences, further supporting the existence of syllable level neural code in the left hemispheric prefrontal regions (Supplementary Fig. 7 & 8). Our results show a clear temporal progression from planning to execution of neural activations associated with the syllable. Further, this neural activation in the planning stage prior to execution supports the idea that syllables function as units of representation for speech planning in left hemispheric prefrontal-planning regions.

Having established the presence of the syllable during speech planning, we next sought to more closely investigate the transition from planning to execution for speech production. One possible process for this transition is that neural populations in prefrontal and sensorimotor brain regions could have distinct activation timing associated with syllable processing during the transition from planning to execution for speech production. These populations with differences in activation timing relative to the utterance onset could form a spatial gradient from planning (prefrontal/anterior) to execution (sensorimotor/posterior) regions and could serve to link these two processes in space and time. This finding would be in keeping with evidence from prior intracranial studies^65^, neuroimaging^11^, and theoretical work^14,16^.

Therefore, to more closely investigate the anatomical and temporal characteristics of the neural populations associated with syllable coding for planning and execution, we used high-channel count, high- density µECoG electrode arrays to record speech neural activations from two subjects. We used two versions of the µECoG arrays to record neural activity along the anterior-posterior orientations from both the planning and the articulation networks (**Fig. 3d**): subject S1, during an awake craniotomy, was acutely implanted with a 256-channel subdural array (12 x 22 matrix; center-to-center electrode distance: 1.72 mm), and subject S2, during an awake deep brain stimulation (DBS) procedure, was acutely implanted with a 128-channel subdural array (8 x 16 matrix; center-to-center electrode distance: 1.33 mm). The implantation procedure for S1 enabled simultaneous recordings from the prefrontal and sensorimotor regions with the array’s longest side oriented parallel to the sylvian fissure. Similarly, the electrode array for S2 was primarily positioned over the prefrontal cortex, with the distal end stretching to sensorimotor cortex. Both subjects completed 4 blocks of the modified speech repetition task. We observed neural activations on individual electrodes with significant modulation in the HG band in both subjects (Supplementary Fig. 9, S1: 201/256 significant electrodes, S2: 63/128 electrodes). Neural activations showed spatially varying temporal characteristics associated with both planning and execution. 18.9% of significant electrodes (Supplementary Fig. 9, S1: 12/201, and S2: 38/63) reached their peak signal power prior to the onset of the utterance. These results indicate the presence of spatially distinct neural population sites that code for speech planning at a micro-scale level recorded with our high-resolution µECoG electrode arrays.

To examine the coding for syllables, we repeated our syllabic discrimination framework analysis on an electrode specific level, by contrasting HG power between disyllabic (VCV) and monosyllabic (CVC) pseudo-word trials. A one-sided permutation test followed by a second-level cluster correction (*p < 0.05*, 1000 shuffles) revealed electrodes that exhibited significant contrasts in activations for the two different syllables, providing further evidence for syllabic coding at the micro-scale (**Fig. 3e**, S1: 100/201, and S2: 37/63). Interestingly, these syllable contrasting electrodes were observed to be spatially organized, and they exhibited distinct temporal profiles identified by onset and peak syllabic coding. For both subjects, we observed spatially varying temporal coding of syllables, with earlier coding starting along the anterior portion of the array and later coding extending towards the posterior side (spatial linear model for S1 using F-statistic test against constant model; Onset coding: F_2,92_ = 6.02, *p = 0.016* and peak coding: F_2,98_ = 10.9, *p < 0.001,* **Fig. 3f**). Comparable prediction results were obtained for S2 where the timings of syllable codes demonstrated a significant spatial anatomical gradient along the anterior-posterior orientation (Peak coding: F_2,28_ = 5.68, *p = 0.024* and Onset coding (F_2,28_ = 6.02, *p = 0.07,* **Fig. 3f**). Taken together, the presence of a spatial anatomical gradient suggests the existence of populations of temporally tuned neurons that activate in succession along the anterior posterior orientation. These populations could form the basis for an anatomical substrate for the transition from planning to execution for speech production.

### 1.4 Two-tiered hierarchical organization for speech production

The ROI-grouped analysis demonstrated how syllables are composed for planning and execution during speech production. However, as syllables are incrementally composed, the speech production system must incorporate appropriate phonemic codes into these structures before generating complete motor articulatory commands during execution^4,13,71^. In other words, the syllable (e.g. CVC) acts as a frame that is filled in with its constituent phonemes (e.g. CVC ◊ /kæt/) We propose that this hierarchical organization is reflected in distinct activation for the syllable and phoneme units in both planning and execution for speech production^14^(**Fig. 4a**).

**Figure 4.**
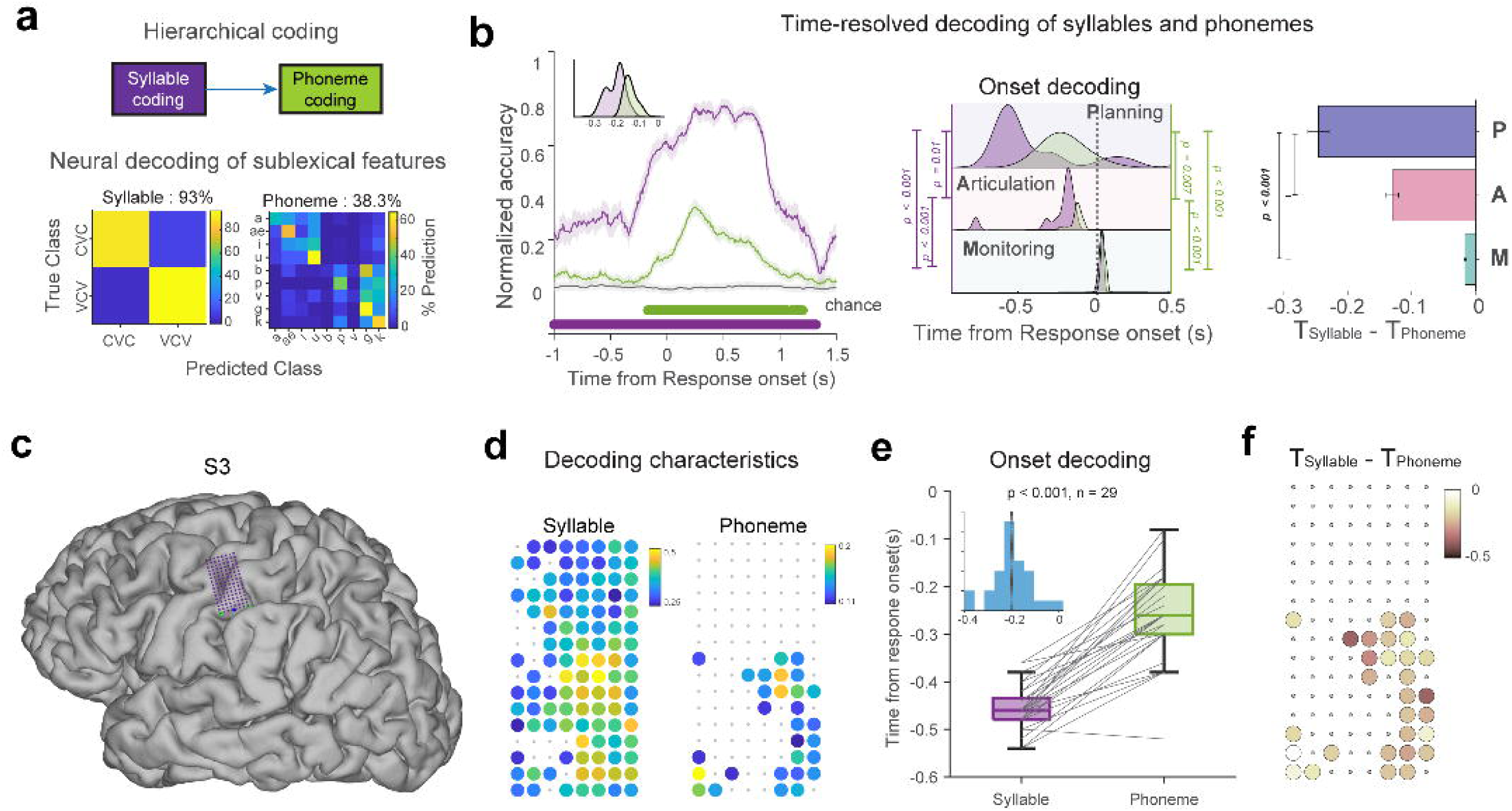
Left hemispheric ROIs demonstrate two-tiered hierarchical organization of syllables and phonemes. **a.** Graphical representation of hierarchical speech representations with syllables coded at higher levels and phonemes are coded at lower levels. The hierarchical organization allows syllable frames to be coded earlier in time, followed by phonemes into the syllable frame. Neural codes for syllables and phonemes were characterized by neural decoding analyses. Heatmaps indicate confusion matrices (SVD-LDA, 10-fold cross-validation) for syllables (2-way; chance 50%) and phonemes (9-way, chance: 11.11%) in speech significant electrodes (2895 total electrodes, 69 mean electrodes per subject). **b.** *Left*: The temporal evolution of syllables and phoneme activations were obtained using time-resolved neural decoding (n = 20, 200 ms window; 10 ms step size). Solid lines indicate average across 20 instances of decoding iterations, and shaded bars indicate standard errors across instances. Grey traces indicate mean and standard errors of chance decoding which was obtained from randomly shuffling syllable and phoneme labels (250 shuffled decoding models). Horizontal bars indicate durations of significant decoding (one-sided temporal permutation cluster test, cluster forming threshold *p < 0.05*; 250 shuffled decoding models). Colored histograms indicate distributions of decoding onsets of syllables (violet) and phonemes (green), with syllables detectable earlier than phonemes (n = 20, one-sided Mann-Whitney U test, *p < 0.001*). *Middle*: Colored histograms indicate distributions of onset decoding times (n = 20) for both syllable and phoneme for each ROI network. Significant differences in decoding distributions indicate a temporal progression across the ROI networks for both syllables and phonemes (n = 20, one-way ANOVA with post-hoc Tukey HSD test, two-sided, *p < 0.01*, for exact p-values see Main text). Syllable coding was earlier in time than phoneme coding (n = 20, *p < 0.05*, one-sided Mann-Whitney U test, for exact p-values see Main text), demonstrating a network-level organization in accordance with the proposed hierarchical representation of syllables and phonemes (see Fig. 1a). *Right*: The difference in activation timing of the hierarchical representation varied for each ROI group, with maximum difference between syllable and phoneme activation during planning (n = 400, 250-350 ms), followed by articulation (n = 400, 110 – 130 ms), and then minimally (n = 400, 17 – 25 ms) during monitoring (n = 400 pairwise differences between syllables and phoneme times per group, one-sided pairwise t-test, planning vs. articulation, onset: t(798) = -10.7, *p < 0.001*, articulation vs. monitoring: onset: t(798) = 18.2, *p < 0.001*; planning vs. monitoring: onset: t(798) = -8.04, *p < 0.001*). Error bars indicate mean differences (± SEM) between syllable and phoneme times for each speech network. **c.** Hierarchical organization at micro-scale population level: µECoG electrode array was implanted over sensorimotor cortex in an awake patient (S3). The electrode array had 128 electrodes with 1.33 mm inter-electrode distance (purple grid). **d**. Electrode-specific decoding characteristics visualized on µECoG grid for syllables (left) and phonemes (right), indicating peak decoding accuracies of electrodes with significant decoding. Syllable decoding was broadly distributed across the array, while phoneme decoding was spatially restricted, indicating a hierarchical topographic organization. **e**. Paired comparison of decoding onset times between syllables and phonemes across electrodes with significant decoding for both features (n = 29 electrodes). Syllable decoding onsets (median = -0.46 s) were consistently earlier than phoneme onsets (median = -0.26s). Box plots show the distribution of decoding onsets for syllables and phonemes (25^th^ – 75^th^ percentile), and the whiskers indicate extreme data points within 1.5 × inter-quartile range of the nearest quartile. The lines connect paired estimates from the same electrodes. Syllable decoding onsets significantly preceded phoneme onsets across the electrodes with a hierarchical lag of 200 ms across the array (T_Syllable – T_Phoneme = -200 ms, one-sided Wilcox signed-rank test, *p < 0.001*). **f**. Spatial map of the onset timing difference across the array. Each circle represents a single electrode with valid syllable and phoneme onset estimates. Color indicates the magnitude of the onset differences, revealing a distributed gradient of hierarchical encoding.

We demonstrated this hierarchical organization by performing temporally resolved neural decoding of speech features (syllables and phonemes) in our speech production tasks from our in-unit patients. We performed a 10-fold, cross-validated decoding (SVD-LDA; see Methods) from HG neural activations (-0.5 s to 0.5 s around utterance onset) using speech significant electrodes across all ROIs pooled across subjects (2895 total grey matter electrodes across both hemispheres, 69 mean electrodes per subject). Significant decoding performance was observed for both syllables (2-way: CVC vs. VCV, average accuracy= 93%, chance = 50%, **Fig. 4a**) and phonemes (9-way: /a/, /æ/, /i/, /u/, /b/, /p/, /v/, /g/, and /k/, average accuracy = 38.3%, chance = 11.11%, **Fig. 4a**). Having established significant decoding for both syllables and phonemes, we sought to characterize the temporal evolution of syllables and phonemes. Using time-resolved decoding (200 ms window; 10 ms step size), we obtained stable decoding for both syllables and phonemes (temporal permutation cluster test, *p < 0.05*, one-sided permutation test against 250 shuffled decoding models). In the left hemisphere, we observed sustained decoding for syllables and phonemes. Crucially, syllables (cluster onset: -1.3s) were detectable at earlier times compared to phonemes (cluster onset: -0.17s, Onset_Syllable_ < Onset_Phoneme,_ one-sided Mann-Whitney U test, *p < 0.001*, **Fig. 4b**), providing evidence that syllables are coded prior to phonemes. We also demonstrated the same temporal order of syllable followed by phoneme within individual ROIs and observed similar stable decoding for both syllables and phonemes (Extended data fig. 3a; except rMFG that exhibited coding only for syllables). For both planning and articulation, we observed syllables represented earlier in time as compared to phonemes (**Fig. 4b**), suggesting that this representational hierarchy is preserved in time both within and across brain areas and speech stages.

Next, we sought to determine whether the temporal relationship between syllables and phonemes is dependent on planning, articulation, and monitoring. We observed a significant difference in onsets of the decoding ability (decoding onsets) for syllables and phonemes from planning to execution, with (as previously mentioned) syllables occurring earlier than phonemes (syllables onset & peak: one-way ANOVA; *p < 0.001*, phonemes onset & peak: one-way ANOVA, *p < 0.001,* **Fig. 4b**, Extended data fig. 3b). Post-hoc Tukey’s HSD test showed that the differences were also pairwise significant (*p < 0.05*) between the three networks, except for peak phoneme decoding between the articulatory and the monitoring networks where the difference did not reach significance (*p = 0.079*). Interestingly, this finding suggests that the magnitude of the difference in timing between the syllable and the phoneme differs as a function of stages/networks.

To assess the change in relative timing of the neural response between syllables and phonemes, we then compared their activation timing within each ROI group. Our results show a maximum delay in the prefrontal-planning network (**Onset**: syllable mean = -0.43s, phoneme mean = -0.22s, difference = 0.25s, *p = 0.039*, one-sided Mann Whitney U test; **Peak**: syllable mean = -0.0125s, phoneme mean = 0.3s, difference = 0.32s, *p < 0.001,* **Fig. 4b**, Extended data fig. 3b), followed by the articulation network (**Onset**: syllable mean = -0.22s, phoneme mean = -0.13s, difference = 0.13s, *p < 0.001*; **Peak**: syllable mean = 0.24s, phoneme mean = 0.35s, difference = 0.11s, *p < 0.001,* **Fig. 4d**, Extended data fig. 3b), and then minimally within the monitoring network (**Onset**: syllable mean = 0.039s, phoneme mean = 0.055, difference = 0.017s, *p = 0.082*; **Peak**: syllable mean = 0.39s, phoneme mean = 0.37s, difference = -0.025s, *p = 0.9,* **Fig. 4b**, Extended data fig. 3b). The mean differences were pairwise significant between all three networks (pairwise t-test, planning vs. articulation, **onset**: t_(798)_ = -10.7, *p < 0.001*, **peak**: t_(798)_ = 71, *p = 0*; articulation vs. monitoring: **onset**: t_(798)_ = 18.2, *p < 0.001*, **peak**: t_(798)_ = 101, *p < 0.001*; planning vs. monitoring: **onset**: t_(798)_ = -8.04, *p < 0.001*, **peak**: t_(798)_ = 117.8, *p < 0.001*, **Fig. 4b**, Extended data fig. 3b). Repeating the analysis with electrodes pooled from right hemispheric ROIs, we observed significant decoding for syllables and phonemes, but we did not observe temporal differences in support of the syllable-phoneme hierarchy (Supplementary Fig. 10).

We further sought to examine this hierarchical organization at the neural-population level. Using a 128-channel µECoG array (1.33 mm inter-electrode distance) over sensorimotor cortex (**Fig. 4c**), we performed electrode specific temporal decoding of syllables and phonemes, separately. Syllable decoding was broadly distributed across the array (108 significant electrodes), while phoneme decoding was spatially restricted to a smaller subset of electrodes (29 significant electrodes, **Fig. 4d**), indicating a topographic organization of the syllable-phoneme hierarchy. Across electrodes with significant decoding for both syllables and phonemes (n = 29 electrodes), syllable decoding onsets (median = -0.46 s; range: -0.48 to -0.435 s) were significantly earlier than phoneme decoding onsets (median = -0.26 s; range: -0.3 to -0.195 s, **Fig. 4e**), corresponding to a median hierarchical lag of approximately 200 ms (T_Syllable_ – T_Phoneme_ = -0.2 s; one-sided Wilcox signed-rank test, *p < 0.001*). Further, the distribution of onset times across the array was entirely negative, confirming a spatially distributed temporal hierarchy at the micro-scale level (**Fig. 4f**).

Overall, these results highlight the hierarchical organization between the syllable and phoneme representation from planning to execution. The differences in hierarchical representation at each speech stage support a transition from separate representations of syllables and phonemes for speech planning in the prefrontal cortex to execution coding in the pre/post central gyrus for articulation and in temporal cortex and insula for monitoring.

### 1.5 Phonological units are sequentially represented only during execution

After planning, the speech production system must successfully articulate the correct ordering of phonological sequences (phonemes) in real-time for the correct execution of the utterance (**Fig. 5a**). To show neural evidence for this sequencing, we performed temporal decoding (9-way; chance 11.11%) to predict the positional phonological units within the pseudo-word utterance in time (e.g. /abæ/: /a/→/b/→/æ/, P1 = /a/, P2 = /b/, P3 = /æ/). A successful sequencing must involve significant decoding of all three phonemes, as well as the correct ordering of phonemes within the sequence. We observed stable decoding and the correct ordering of sequences (P1→P2→P3) in the ROIs specific to the articulation and monitoring networks. Surprisingly, this ordering was absent (absence of phoneme positions or incorrect ordering) in the planning network ROIs (p < 0.05, one-sided permutation test against 250 shuffled decoding models, **Fig. 5b**), indicating that phoneme codes retrieved during planning are not explicitly sequenced at this stage, rather, they are sequenced in real-time in the motor cortex. In the articulation ROIs, we observed pairwise significant differences between phoneme positions in peak decoding times (T_P1_ < T_P2_ < T_P3_; one-sided Mann Whitney U test, *p* < 0.05; T_P1_ < T_P2,_ *p < 0.001*; T_P2_ < T_P3_, *p < 0.001*. T_P1_ < T_P3_, *p < 0.001,* **Fig. 5a**). The monitoring ROIs exhibited similar positional phoneme coding in peak times (T_P1_ < T_P2_ < T_P3_; one-sided Mann Whitney U test, *p* < 0.05; T_P1_ < T_P2,_ *p < 0.001*; T_P2_ < T_P3_, *p < 0.001*. T_P1_ < T_P3_, *p < 0.001,* **Fig. 5a**), which demonstrate continuous monitoring from auditory feedback. Similar ordering of phonological sequencing was also observed in the articulation and the monitoring network in the right hemisphere (Supplementary Fig. 11). Taken together, the results show that for sequencing, only the articulation network generates positional phonological unit codes in their correct order. This is carried out during the execution of speech, along with the monitoring network which examines the phonological sequence of the utterance through feedback.

**Figure 5.**
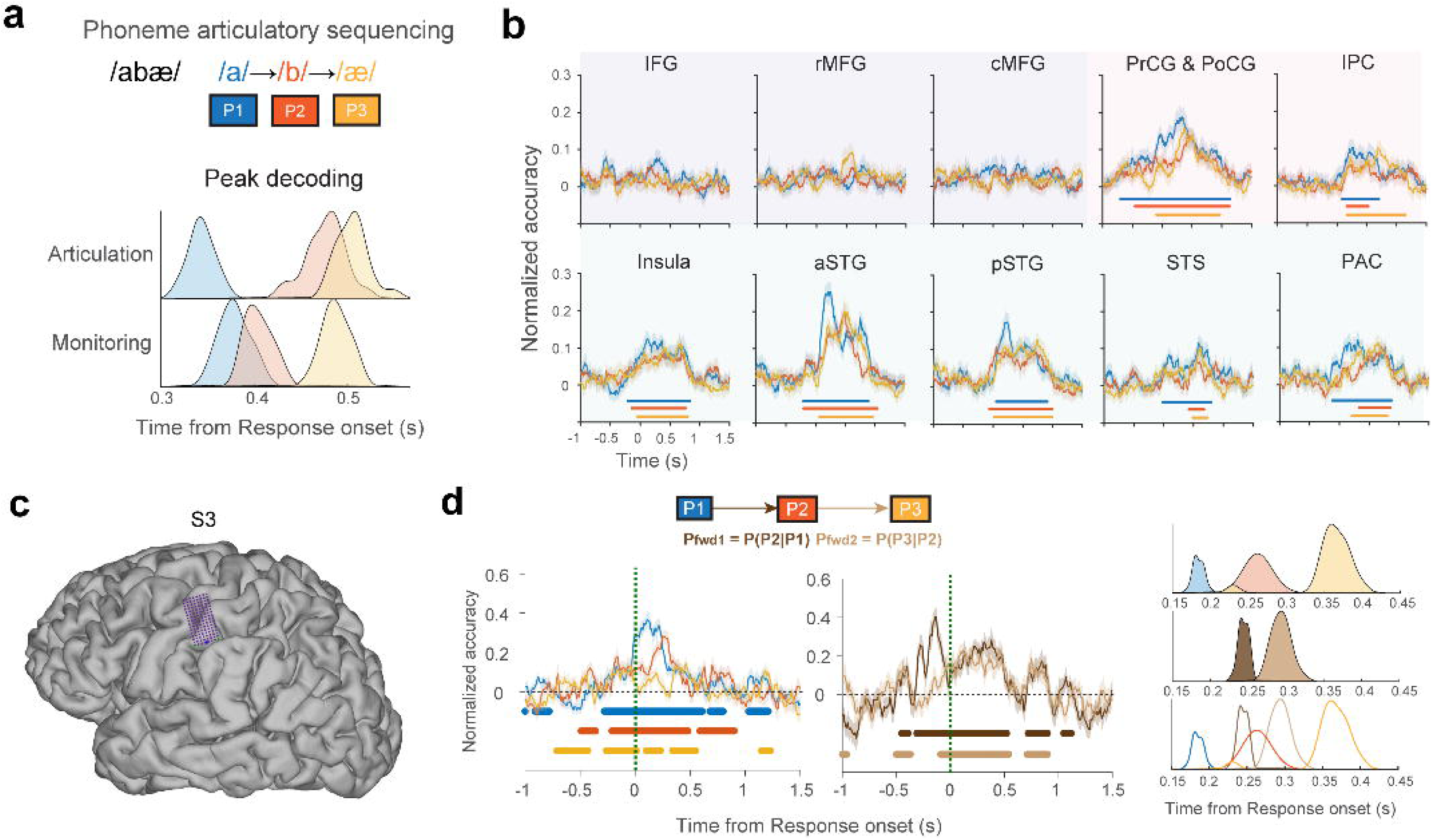
Phonological units are sequentially articulated during execution. **a.** Graphical representation of sequential execution that depicts phonological units articulated in their correct order during the utterance. Colored histograms show distribution times of decoding peaks for each positional phonological unit within the ROI network. Positional phonological unit codes were decoded in the correct serial order (pairwise one-sided Mann-Whitney U test, *p < 0.05,* for exact p-values see Main text) **b.** Time-resolved decoding models were trained on neural signals aligned to the utterance to decode phonological units at each position within the pseudo-word. Each colored trace indicates temporal evolution of positional phonological units in time for a ROI (mean across 20 decoding instances). Results are shown for all ROIs in the left hemisphere, grouped by planning, articulation, and monitoring networks. Significant phonological unit decoding for all positions was observed only in execution ROIs (articulation and monitoring), suggesting that temporal sequencing is carried out as part of the execution of speech sounds rather than during planning. Horizontal bars indicate durations of significant decoding (one-sided temporal permutation cluster test, cluster forming threshold *p < 0.05*; 250 shuffled decoders). Background shading indicates ROI networks for planning, articulation and monitoring. The presence of ordering demonstrates the serial execution of positional phonological unit codes by the execution network. **c**. Continuous phonotactic transitions are present in motor cortex during articulation. µECoG electrode array was implanted over sensorimotor cortex in an awake patient. The electrode array had 128 electrodes with 1.33 mm inter-electrode distance (purple) **d.** Graphical schematic demonstrating that phonological unit sequence execution may involve both discrete phonological units as well as their underlying transitionary statistics (phonotactics). Forward phonotactic transition from C1 to V2 is denoted by P_fwd1_, and from V2 to C3 is denoted by P_fwd2_. Time-resoled decoding revealed accurate phonological unit sequencing for CVC trials (C1: blue, V2: orange, C3: yellow). Temporal decoding models trained to predict the phonotactic transitions (3-way: low, medium, and high phonotactic transitions) demonstrated significant decoding for P_fwd1_ (dark brown) and P_fwd2_ (light brown). Horizontal bars indicate durations of significant decoding (one-sided temporal permutation cluster test, cluster forming threshold *p < 0.05*; 250 shuffled decoders). Colored histograms indicate distribution times of peak decoding onsets for positional phonological units and phonotactic transitions, revealing significant temporal ordering of phonological units and phonotactic transitions (pairwise one-sided Mann-Whitney U test, p < 0.05, T_C1_ < T_V2,_ *p < 0.001*; T_V2_ < T_C3_ *p < 0.001*, T_Pfwd1_ < T_Pfwd2,_ *p < 0.001*). Further, the timing of the forward transitions was embedded between the phonological unit positions (pairwise one-sided Mann-Whitney U test, p < 0.05, T_C1_ < T_Pfwd1,_ *p = 3.7e-08*; T_Pfwd1_ < T_V2_, *p < 0.001*, T_V2_ < T_Pfwd2_, *p < 0.001*, T_Pfwd2_ < T_C3_, *p < 0.001*). This significant temporal embedding demonstrates the existence of both discrete and continuous motor coding during the execution of sequences during speech production.

Lastly, we investigated the specific neural processes involved in speech sequencing during articulation. While we showed that activation of discrete ordered phonemes was present during execution, this activation could reflect discrete motor plans for each phoneme, or sequences of continuous motor activation that reflect the execution of the entire sequence. If the entire sequence is executed holistically, then we should see evidence for both discrete and continuous motor processes during articulation.

To demonstrate both discrete and continuous processing, we examined neural activation associated with the transitions between phonemes in the sequence. These transitions reflect the statistical properties of the distribution of phoneme co-occurrences from the word patterns specific to the native language (English in this case). We posited that the presence of information about transitions between phonemes would reflect evidence for continuous rather than discrete processing of the utterance^72,73^(**Fig. 5d**).

We recorded micro-scale cortical activations in one subject (S3) using the 128-channel high-density µECoG electrode array over speech motor cortex (SMC) (**Fig. 5c**). We found significant HG activations associated with speech production (111/128 electrodes, see also patient S1 in our previous work^63^). To assess whether the transitions between phonemes were tracked in the SMC, we focused on trials with monosyllabic structures (CVC; to remove the confound of two syllables), by measuring the ability to decode the forward phonotactic transitions (P_fwd1_: P(P2|P1) & P_fwd2_: P(P3|P2)). The forward transitions were estimated by calculating the conditional probability of one phoneme given the previous, based on positional and co-occurrence counts. The resultant transition probabilities were quantized into three evenly distributed classes producing trials with low, medium, and high transitions scores (P_fwd1_: low – 0.0024 to 0.034, medium – 0.034 to 0.1031, high – 0.1031 to 0.1791; P_fwd2_: low – 0.0083 to 0.0314, medium – 0.0314 to 0.0543, high – 0.0543 to 0.0814).

Our time resolved decoding again showed evidence of the correct order of phonological units (C: 5-way, V: 4-way) within the CVC trials (P1→P2→P3, **Fig. 5d**). Next, we repeated a similar decoding analysis to predict the forward transition scores (3-way) and computed these values in time. We found that the decoding for P_fwd1_ occurred earlier in time compared to P_fwd2._ Within features, we observed significant temporal ordering of phonological units (T_P1_ < T_P2_ < T_P3_; one-sided Mann Whitney U test, *p* < 0.05; T_P1_ < T_P2,_ *p < 0.001*; T_P2_ < T_P3_, *p < 0.001*, **Fig. 5d**) and phonotactic transitions (T_Pfwd1_ < T_Pfwd2,_ *p* = 6.6e-08). Across features, the forward transitions were significantly embedded between the phonological unit positions (T_P1_ < T_Pfwd1_ < T_P2_ < T_Pfwd2_ < T_P3_; pairwise one-sided Mann-Whitney U test, *p < 0.05*; T_P1_ < T_Pfwd1,_ *p < 0.001*; T _Pfwd1_ < T_P2_, *p < 0.001*, T_P2_ < T_Pfwd2_, *p < 0.001*, T_Pfwd2_ < T_P3_, *p < 0.001*, **Fig. 5d**). These results indicate that the neural basis of speech sequential processing during articulation reflects both discrete activation of phonological units, as well as the continuous processing of the phonotactic transition between these units.

## 2 Discussion

Speech planning requires the brain to encode the structural and phonemic content of intended utterances and organize them into sequences before motor execution begins. Prior neuroimaging work implicated prefrontal regions in this process, but their precise functional role and how planning relates temporally to execution and monitoring remained underspecified. Using intracranial recordings during pseudo-word repetition, we characterized the neural substrates of speech planning and execution at the sub-lexical level, and provide direct electrophysiological evidence for a hierarchical, cascade-like organization of speech production^11,13^. Together, these findings establish a framework for understanding how the brain transitions from planning abstract speech units to executing them as continuous sequences.

We identified three networks distinguished by the sequential timing of their high-gamma activations, providing a direct measure of the temporal organization of speech production. Planning activations in prefrontal regions (IFG, rMFG, cMFG) preceded articulation regions (precentral/postcentral gyrus, inferior parietal cortex) by a median of 180 ms, which in turn preceded auditory-monitoring regions (STG, STS, insula) by 398 ms, and finally a 370 ms planning-to-monitoring lag. This ordering was independently confirmed at the subject level through functional connectivity analysis and replicated by unsupervised decomposition, which recovered the same three networks without any prior anatomical constraints. The role of IFG and MFG has historically been framed around phonological working memory and sub-lexical retrieval^65^, but our data reframe this: prefrontal regions encode syllabic structure prior to articulation by sensorimotor and inferior parietal regions^14,20,58,74^, pointing to a role in organizing sub-lexical representations for execution. Late activations in STG and STS were consistent with auditory monitoring and feedback prediction, though two temporally distinct auditory components were identified, suggesting that monitoring has more complex dynamics than a single feedback loop^75,76^. Taken together, the cascade-like timing across these three networks supports a largely feedforward flow of information and argues against interactive-parallel accounts of speech production.

The hierarchical relationship between syllables and phonemes is a central prediction of DIVA, GODIVA, and Hickok’s HSFC models, each of which proposes that higher-level planning states code syllabic frames while lower-level states handle phonemic content^13,14,16,27^. We provide direct electrophysiological support for this view, and critically show that this hierarchy compresses systematically as information flows from planning to execution. In planning regions, the syllable-to-phoneme lag was 250–350 ms; in articulation regions, this compressed to 110–130 ms; and in monitoring regions, the two representations were nearly concurrent, separated by only 17–25 ms. The compression window closely matches the 180 ms planning-to-articulation connectivity delay, suggesting that inter-network communication itself may be the mechanism through which hierarchical structure is resolved into sequential motor commands. At the micro-scale, electrode-specific decoding in sensorimotor cortex confirmed syllable-first ordering with a median lag of 200 ms, topographically restricted to the ventral portion of the array, indicating that temporal compression is a spatially localized computational property of sensorimotor cortex. This systematic compression across stages represents a novel temporal finding that existing models do not account for and that future computational frameworks will need to incorporate^14^.

Sequential phoneme encoding is a fundamental requirement for fluent speech and understanding where and how it emerges in the production network is critical for both basic science and clinical applications^13^. While syllabic coding was robust in planning regions, sequential decoding of phoneme order was absent there and emerged only in articulation and monitoring networks, suggesting that phonemes are dynamically sequenced during motor execution rather than pre-ordered during planning^58,66,74,77^. Beyond discrete phoneme sequencing, sensorimotor cortex also encoded phonotactic transition probabilities^73^, with activations emerging temporally between individual phoneme position activations, indicating that motor articulation is organized as a continuous process shaped by internalized phonological statistics. This finding has direct relevance for speech BCI design^78–81^, where models that capture only phoneme identity will be insufficient to reconstruct the continuous, statistically structured motor dynamics that underlie fluent articulation.

While this study provides high-resolution temporal maps of sub-lexical planning and execution, several important limitations point to directions for future work. Our findings are correlational, and the directional information flow implied by connectivity timing must be tested causally through perturbation protocols or stimulation paradigms. The contribution of working memory to the maintenance of sub-lexical representations during planning remains uncharacterized in this paradigm, and future studies will examine how prefrontal working memory capacity interacts with the planning of syllabic and phonemic content prior to articulation. In addition, extending the paradigm to include stimuli with varying phonological and phonotactic properties will determine how sub-lexical features shape the planning-to-execution pipeline, and introducing lexical properties will further connect this pipeline to language production more broadly, bridging the gap between sub-lexical motor encoding and higher-order linguistic processing^4^.

The timing architecture described here, including cascade onset delays, syllable-phoneme compression gradients, and spatially organized motor sequencing, provides a concrete empirical target for updating next-generation computational models of speech production. More broadly, characterizing the spatio-temporal organization of sub-lexical planning and execution has direct implications for the development of speech neuroprostheses and for understanding how disruptions to these networks give rise to speech and language disorders.

## 3 Methods

This study was performed at the Duke University Health System with approval from the Institutional Review Board under the protocol IDs:

Pro00065476: Network dynamics of human cortical processing.

Pro00072892: Studying human cognition and neurological disorders using µECoG electrodes.

### 3.1 Participants

Intracranial recordings with standard clinical intracranial recording systems were obtained from 52 patients with pharmaco-resistant epilepsy at Duke University Medical Center (DUMC) (mean age = 33.1 years, 23 Female, Supplementary table 1). Patients were implanted with clinical intracranial electro-encephalography (IEEG) systems as part of their work up for the neurosurgical treatment of pharmacologically resistant epilepsy. Recordings using high-density micro-scale cortical recordings were obtained in an intraoperative setting from 3 adult patients (mean age = 64.3 years, 0 Female), during awake neuro-surgical procedures. Patients were awoken as part of their surgical procedure and the decision to have them awake was entirely clinically motivated. Two of the three patients (S1 and S3) underwent the placement of a deep brain stimulator (DBS) for the treatment of movement disorders. The other patient (S2) underwent surgical resection for tumor removal. All patients were native English speakers and written informed consent was obtained in accordance with the Institutional Review Boards at the DUMC.

### 3.2 Electrode implantation, localization and neural recordings

Cortical recordings were obtained using electrodes from either one of the three recording systems: 1) macro-electrocorticographic grids (macro-ECoG: 2 participants), 2) stereo-electroencephalographic probes (SEEG: 50 participants), and 3) liquid crystal polymer thin-film (LCP-TF) micro-electrocorticographic arrays (µECoG: 3 participants). Macro-ECoG grid recordings were done using Ad-Tech electrode grids (48 or 64 electrodes; Ad-Tech Medical Instrument Corporation) with 10 mm electrode spacing and 2.3 mm exposed diameter, surgically implanted onto the cortex of the patients. SEEG placement was done via robotically guided surgery (ROSA Robotics) using either PMT (PMT Corporation), Ad-Tech, or Dixi (Dixi Medical) SEEG electrodes with 3.5 mm to 5 mm inter-electrode spacing (0.8 to 0.86 mm exposed diameter). µECoG recording included LCP-TF µECoG arrays (1.33 – 1.72 mm electrode spacing; 200 µm diameter), acutely implanted either through a burr hole (DBS surgery) or directly over the cortical surface (awake craniotomy).

Pre-operative anatomical 3T MRI were obtained for all patients, and post-operative CT images were obtained for both macro IEEG and DBS patients to identify electrode locations. Patients undergoing DBS surgery underwent a routine intraoperative CT for assessment of DBS electrode placement with the subdural ECoG array in place. For the tumor resection patient, BrainLab coordinates^82,83^ were obtained at the corner of the electrode array and this information was then mapped to three-dimensional reconstruction of the patient’s brain obtained from their preoperative MRI. For each patient, electrode locations were manually marked using BioImage Suite, and were mapped on to the reconstructed T1 volume generated using Freesurfer^84,85^. Electrodes were then assigned to specific regions of interest (ROIs), based on the cortical parcellations obtained from connectivity based Brainnetome Atlas^86^. For the electrodes that were not centered at specific ROIs (e.g. depth electrodes from SEEG probes near/in white matter), a sphere with a 10 mm radius was constructed around the electrode’s center, and the relative density of ROIs was determined through the calculation of the percentage proportion of ROI voxels within this sphere. Electrodes were then assigned to a specific ROI with the maximum density if the proportion exceeded 5% (otherwise they were labelled as white matter contacts). For visualization, electrode coordinates were also transformed to a population averaged MNI brain and color-coded for ROIs.

IEEG recordings in the clinical monitoring unit (ECoG and SEEG) were analog filtered between 0.01 Hz and 1000Hz and digitized at 2,048 samples per second, using a Natus Quantum LTM amplifier (with Natus Neuroworks EEG software). Neural recordings from µECoG electrodes (intra-operative) were analog filtered between 0.1 Hz and 7500 Hz and digitized at 20,000 samples per second (with Intan RHX data acquisition software, v3.0; Intan Technologies, Inc).

### 3.3 Delayed speech repetition task

Patients performed a delayed pseudo-word repetition task in the clinical monitoring unit. In each trial, they listened and repeated one of 52 pseudo-words that were modelled on the phonology of American English. Pseudo-words were equally distributed between monosyllabic and di-syllabic structures (consonant-vowel-consonant - CVC: 26) and vowel-consonant-vowel (VCV: 26 tokens) and included a total of 9 phonemes (4 vowels: /a/, /æ/, /i/, /u/ and 5 consonants: /b/, /p/, /v/, /g/, /k/). The phonemes were arranged within the pseudo-words (CVC or VCV) by selecting biphones (CV or VC) with a positive likelihood of occurring in the English language corpus^87^. The third phoneme was selected with the constraint that the occurrence probability of the triphone sequence was zero, resulting in a constructed pseudo-word. For each pseudoword, forward biphoneme transition probabilities were calculated based on the occurrence probabilities of individual phonemes [P(C) or P(V)] and co-occurring biphones [P(CV) and P(VC)] as follows:

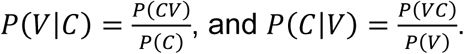

The resultant transitional probability scores across the pseudoword tokens were quantized into three classes: low, medium, and high transitions scores (P(V|C): low – 0.0024 to 0.034, medium – 0.034 to 0.1031, high – 0.1031 to 0.1791; P(C|V): low – 0.0083 to 0.0314, medium – 0.0314 to 0.0543, high –0.0543 to 0.0814).

The constructed pseudo-words were auditorily presented and subsequently cued for the patients to repeat via a visual Go Cue (‘speak’). Each trial lasted between 5.1 – 6.1 seconds, starting with a visual “Listen” cue followed by the auditory stimulus presentation (mean stimulus duration: 0.5 second). After a 1.5 s delay (150 ms jitter), the “speak” (Go) cue was visually presented and the patients were given 3 s (200 ms jitter) to respond, before the next trial began. Trials were presented in four consecutive blocks with 52 unique pseudo-word tokens shuffled in each block and the overall task time with 208 speech trials took less than 40 minutes in the monitoring unit. A subset of patients underwent a similar delayed repetition task where they repeated pseudo-words with di-syllabic structures (CVCVC). The patients in the intra-operative setting underwent a similar version of pseudo-word repetition task, but with the delay and “speak” cue removed for simplicity. Intraoperative patients completed 3 to 4 blocks of 52 trials (156 – 208 trials). The speech task was designed using Psychtoolbox scripts in MATLAB R2014a.

### 3.4 Audio recording and behavioral representation

The speech repetition task included both auditory and speaking components. The auditory stimulus wave files were presented using a laptop connected to a powered stereo-speaker (amazonbasics A100 series) through a USB DAC and audio amplifier (Fiio). The beginning of each trial and the subsequent stimulus onsets were marked by flashing white circles on the stimulus laptop screen which was detected by a photodiode (Thorlabs, Inc.) attached to an auxiliary BNC-to-mono TS cable connected to the Natus Recording system. The audio waveforms were recorded using a condenser microphone (Marantz Professional, MPM-1000) which was connected to pre-amplifier (Behringer) and digitized at 44100 Hz on the recording laptop. The auditory stimulus, the spoken pseudo-word onsets, and the offsets were either manually identified using the Audacity software package, with the built-in Mel-spectrogram visualization feature or automated through the Montreal Forced Aligner (Montreal Corpus Tools).

For behavior analysis, each trial was characterized by its response time (with respect to Go cue onset) and its response duration. For the neural analysis, the trials were categorically coded for their syllabic structure (CVC or VCV), the phonemes (9-way) at each position, and the corresponding forward transition category (3-way) for each biphoneme pair within its structure. Trials with no intelligible responses were removed from the analysis.

### 3.5 Behavioral representations of syllabic coding

A linear mixed effects (LME) model was used to predict the effects of syllable on response time and response duration. The models incorporated fixed effects for categorical syllabic IDs (one vs. two syllables) and a random intercept for patient. Two sets of models were utilized separately to predict response time and response duration respectively, and in each case the estimate of syllable (beta coefficient for fixed effect) was computed. The significance of the beta estimates was computed using a permutation test by randomly shuffling syllables (1000 shuffles) and repeating the LME prediction for each shuffle.

### 3.6 Neural preprocessing and signal analysis

The neural preprocessing steps were performed offline. The IEEG recordings were filtered for line-noise with multi-taper band-stop notches centered at 60, 120, 180, and 240 Hz. The following categories of electrodes were removed from subsequent analyses: 1) electrodes that were outside the cortex (determined from localization), 2) electrodes that exhibited muscle artifacts during the utterance (as visualized by a wavelet scalogram), and 3) electrodes with recording power greater than 3 * log-RMS. From a total of 9679 electrodes, 1573 were excluded using the above steps. The remaining set of 8106 electrodes were subjected to common-average-referencing (CAR) and utilized for further processing.

#### 3.6.1 Neural feature extraction

Analyses were performed to extract neural high-gamma signals (HG: 70 – 150 Hz) from each electrode. HG signals have previously been shown to characterize speech activations from cortical regions as well as demonstrating a high correlation with multi-unit firing^54,56,57^. The pre-processed neural signals from each IEEG electrode were band-pass filtered into 8 center frequencies (logarithmically arranged) between 70 and 150 Hz^88^. The envelope of each frequency band was extracted using the Hilbert transform and was averaged across the bands. The resultant envelope was then downsampled to 200 Hz using an anti-aliasing filter and z-scored with respect to a baseline period (-500 ms to 0 s before trial start). For each trial, we defined the response epoch as the time intervals from -1500 ms to 1500 ms with respect to the utterance onset.

#### 3.6.2 Functional connectivity analysis

To assess the temporal ordering of speech networks at the subject level, pairwise net cross-correlations were computed between the speech networks: planning, articulation, and monitoring, for each subject that contributed electrodes to atleast two networks. Random subsets of up to 50 electrodes were sampled from a network-pair through 500 bootstrap iterations, and the normalized cross-correlation was computed between each sampled electrode pair in the high gamma band. To isolate between-network coupling from shared within-network correlation structure, a net cross-correlation was derived by subtracting the average of the two within-network autocorrelations from the between-network cross-correlation. The peak lag of the net cross-correlation was identified per electrode pair, and the median peak lag was obtained per subject per network pair. To test the directional significance of temporal structure, a circular-shift null distribution was constructed by applying a random time-shift offset to one electrode signal prior to cross-correlation, thereby preserving the autocorrelation structure while randomly modifying the between-network temporal structure, and the significance was calculated using a one-sided paired Wilcox signed-rank test.

#### 3.6.3 Non-negative matrix factorization

To perform a data-driven quantification of neural sub-components underlying speech production, we performed non-negative matrix factorization (NNMF) on the neural HG power^89^. HG activations (***X***) were pooled across all speech-responsive electrodes and were averaged across the trials with respect to the utterance onset (from -1.5 seconds to +1.5 seconds). NNMF was used to decompose the electrode-time matrix into two non-negative matrices, *X* ≈ *H* × *W*, where ***H*** contains the electrode weights (channels x components) and ***W*** captures the temporal profiles of each component (components x timepoints). Factorization was performed using an alternating least-squares algorithm with non-negativity constraints on both ***H*** and ***W***. The optimal number of components (k) was determined by evaluating the cumulative variance explained at each candidate k and selecting the factor that accounted for at least 90% of the data variance.

#### 3.6.4 Neural representations of syllable coding

A non-parametric statistical test was performed to test whether HG power from each ROI could differentiate between syllables (CVC vs. VCV) at each time-point. For each electrode, the absolute difference in HG power (ΔHG) was computed between VCV trials and CVC trials. The resultant difference (across-syllables) was averaged across electrodes within a given ROI (within patient followed by across patient average) and was considered as the test statistic. For a control analysis, ΔHG was computed within CVC trials and within VCV trials, and the average within-syllable ΔHG was determined for each ROI. The null distribution was generated from 1000 iterations of the within-syllable statistic for each ROI, and the significance was determined by calculating the proportion of null-distribution (within-syllable) less than the test-statistic (across-syllable) (*p < 0.05*). The resultant p-value time-series were subjected to a second-level non-parametric cluster correction to determine consecutive sequence of time-points that exhibited significant syllable coding.

To rule out the confounding of initial phonemes (C vs. V) in the neural syllable code, a control analysis was performed in a subset of patients who underwent a modified version of speech repetition task, where they repeated pseudowords with di-syllabic structures (CVCVC). Similar to the above method, the absolute difference in HG power was computed between CVCVC trials and CVC trials as the test statistic and was subjected to second-level non-parametric cluster correction.

#### 3.6.5 Linear classification model for speech decoding

A previously established linear-classification model with 10-fold cross-validation, was used to decode speech units from normalized HG^63^. First, we performed a singular value decomposition (SVD) to compress HG to low-dimensional features. We chose the number of kept components as the total number of components that explained 80% of the variance. We utilized a supervised linear-discriminant (LDA) model to predict speech units. The decoding performance was estimated by calculating the average prediction accuracy in the test set across all folds.

Multivariate decoding was performed by pooling HG epochs from electrodes across patients. To perform this pooling, trials with similar tokens across patients were concatenated together. In instances where there was a mismatch in the number of trials across patients for a particular token, a synthetic trial was generated using a Mix-Up technique through the linear combination of two randomly selected trials for that token^90^. This Mix-Up approach was performed separately on the training and the testing set to avoid data leakage.

To infer temporal characteristics of speech features, decoding was performed in the temporal domain by training and testing at each time-segment (200 ms window; 10 ms step size) within the epoch^91^. The decoding accuracy was calculated at each time-segment to characterize temporally evolving representations of speech features in the response epoch. Separate temporal models were evaluated per ROI to decode syllables (2-way, chance: 50%), phoneme units (9-way, chance: 11.11%) in each position (P1, P2, and P3), and phoneme transition probabilities (3-way, chance: 33.33%) for each biphone (P_fwd1_, and P_fwd2_) within the pseudo-word. The resultant accuracy (Acc_Test_), as a percentage, was normalized (Acc_Norm_) with respect to theoretical chance (Acc_Chance_) for visualization purposes as follows:

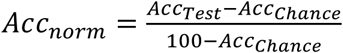

#### 3.6.6 Statistical analysis

Multiple statistical tests were conducted to validate IEEG recordings for speech representations. To identify electrodes with significant speech-neural activations, signal power in the HG band was compared between the response (-500 ms to 500 ms) and the baseline period using a 1000-iteration one-sided permutation test (corrected for false discovery rate).

To quantify the neural morphology of HG responses for each ROI, the above non-parametric test was extended to determine the significance at each time point (against baseline), with HG power averaged across electrodes within the ROI. In addition, the HG onset times, and peak times were calculated for each electrode. The onset times were estimated by performing a change-point analysis which determines the time-point at which the mean of the HG time-series changes most significantly (findchangepts algorithm in MATLAB). Peak times were determined by either finding the time-point where HG activations achieved the maximum value or by computing the weighted mean of HG activations over time. The distribution of HG onset and peak times were used to quantify the evoked temporal characteristics of a speech responsive electrode. Significant differences in onset or peak times were evaluated using pairwise Mann-Whitney U test, or student t-tests.

For speech decoding, the prediction accuracies were validated against the null distribution obtained from random shuffling of the predictor labels (250 iteration). ROI characteristic decoding onsets were obtained using a similar change-point analysis. The resultant time-series was subjected to second-level non-parametric temporal cluster correction.

## Data availability

Derived data supporting the analyses reported here are openly available in the Zenodo repository (https://doi.org/10.5281/zenodo.20720848). These permit reproduction of the analyses reported in this paper.

Raw iEEG recordings are not publicly available because they constitute potentially identifiable human neural data and the study’s current IRB protocol and participant consent do not cover open dissemination. De-identified raw recordings, which support independent re-processing and analyses beyond those reported here, are available to academic researchers for non-commercial research purposes under a data use agreement specifying permitted analyses and prohibiting re-identification or redistribution. Requests should be directed to the corresponding authors and will be answered within 2 weeks.

## Code availability

The MATLAB files to perform neural analysis is available through GitHub at: https://github.com/coganlab/IEEG_Pipelines & https://github.com/coganlab/speechPlanning_manuscript.

Claude Sonnet models were used for generating visualization scripts and for documentation purposes. All LLM generated code was verified by the researchers.

## Acknowledgements

We would like to thank Anna Thirakul for help with consenting and IRB compliance, Palee Abeyta, Aaron Earle-Richardson, and Nicole Liddle for help with data collection for the in-unit patients with epilepsy, Seth Foster and Aaron Earle-Richardson for helping with visualization software and data pipelining, and James Carter for help with intraoperative data collection. We would also like to thank the patients who graciously volunteered their time for this study.

## Author Contributions Statement

S.D. performed neural feature analysis, developed neural decoding models, performed statistical assessments, and generated figures. G.B.C., and S.R. designed the experimental task. S.R., S.C.H., D.P.S., S.P.L., A.H.F., and D.G.S., were involved in surgical planning and performed intra-operative electrode implantation and neural data acquisition. K.B., C.H.C., C.W., and J.V. were involved in µECoG electrode design and setting up intra-operative data acquisition process. D.G.S., and S.R.S assisted in extra-operative data acquisition from SEEG and ECoG electrodes. S.R., J.V., and G.B.C contributed to neural feature analysis and the development of speech decoding models. G.H. provided conceptual guidance for the manuscript. S.D., J.V., and G.B.C., wrote the manuscript with feedback from the other co-authors. G.B.C. was involved in clinical supervision. G.B.C., supervised the human in-unit monitoring study, and J.V., and G.B.C. designed and supervised the human intra-operative study.

---

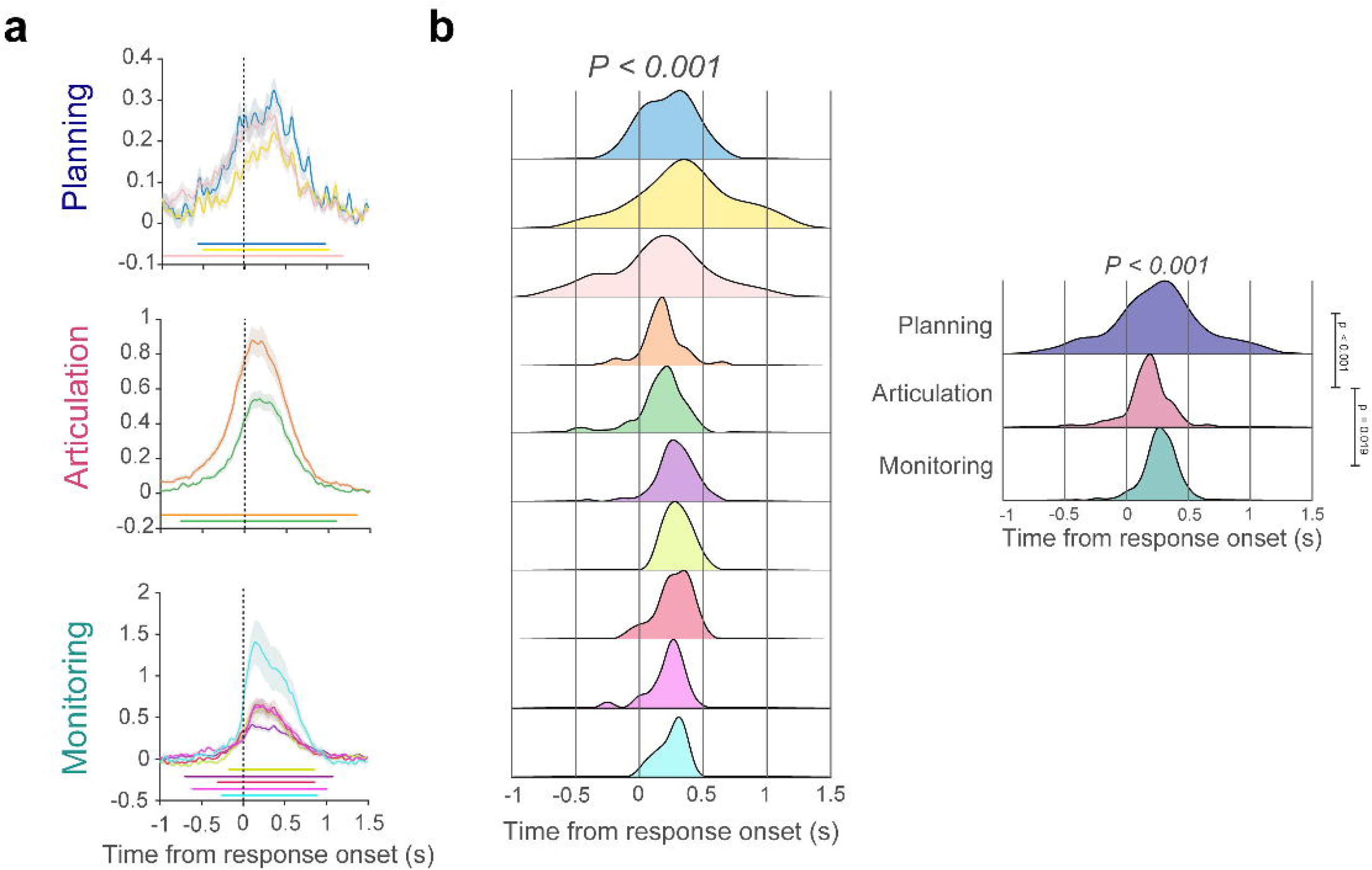

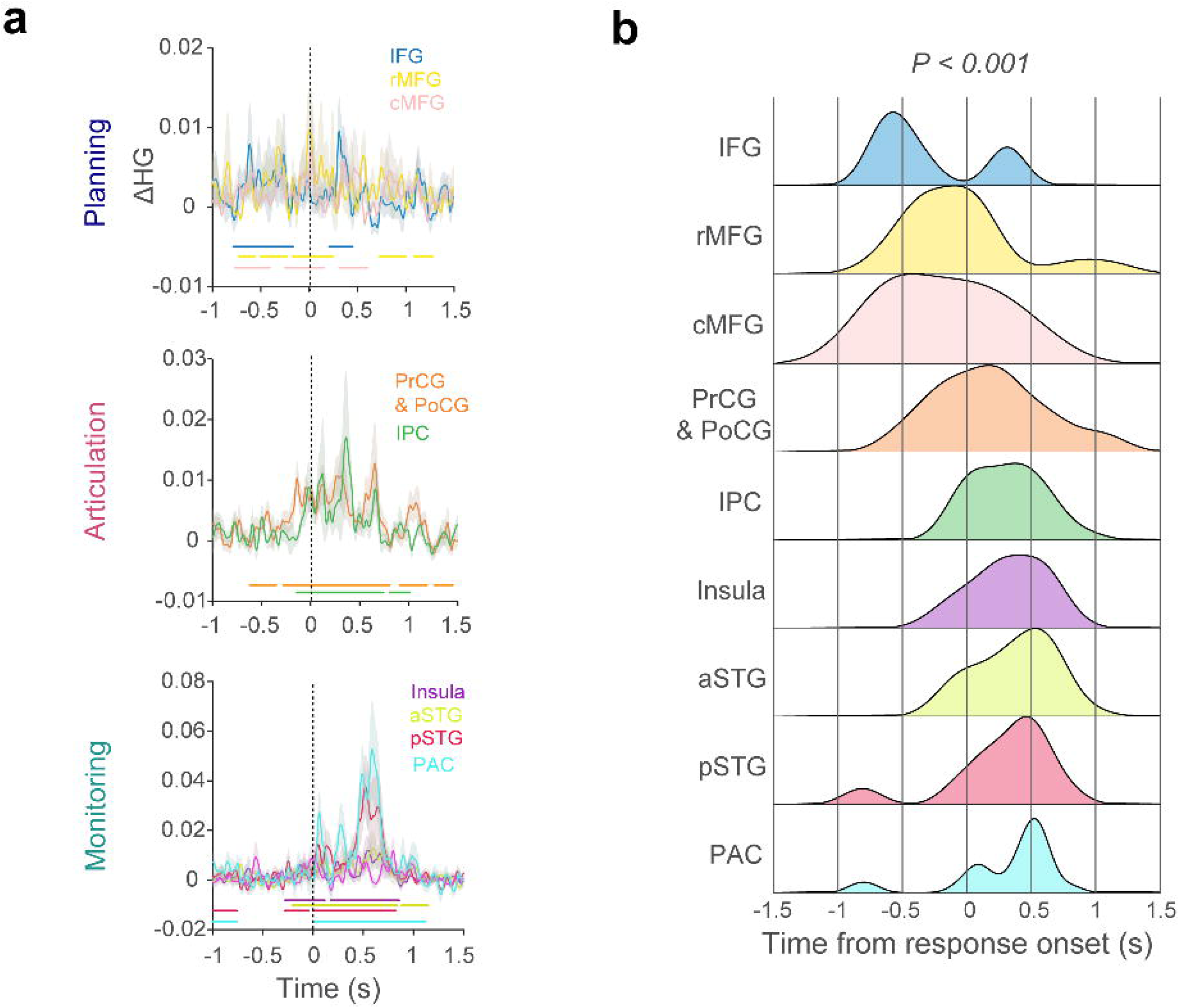

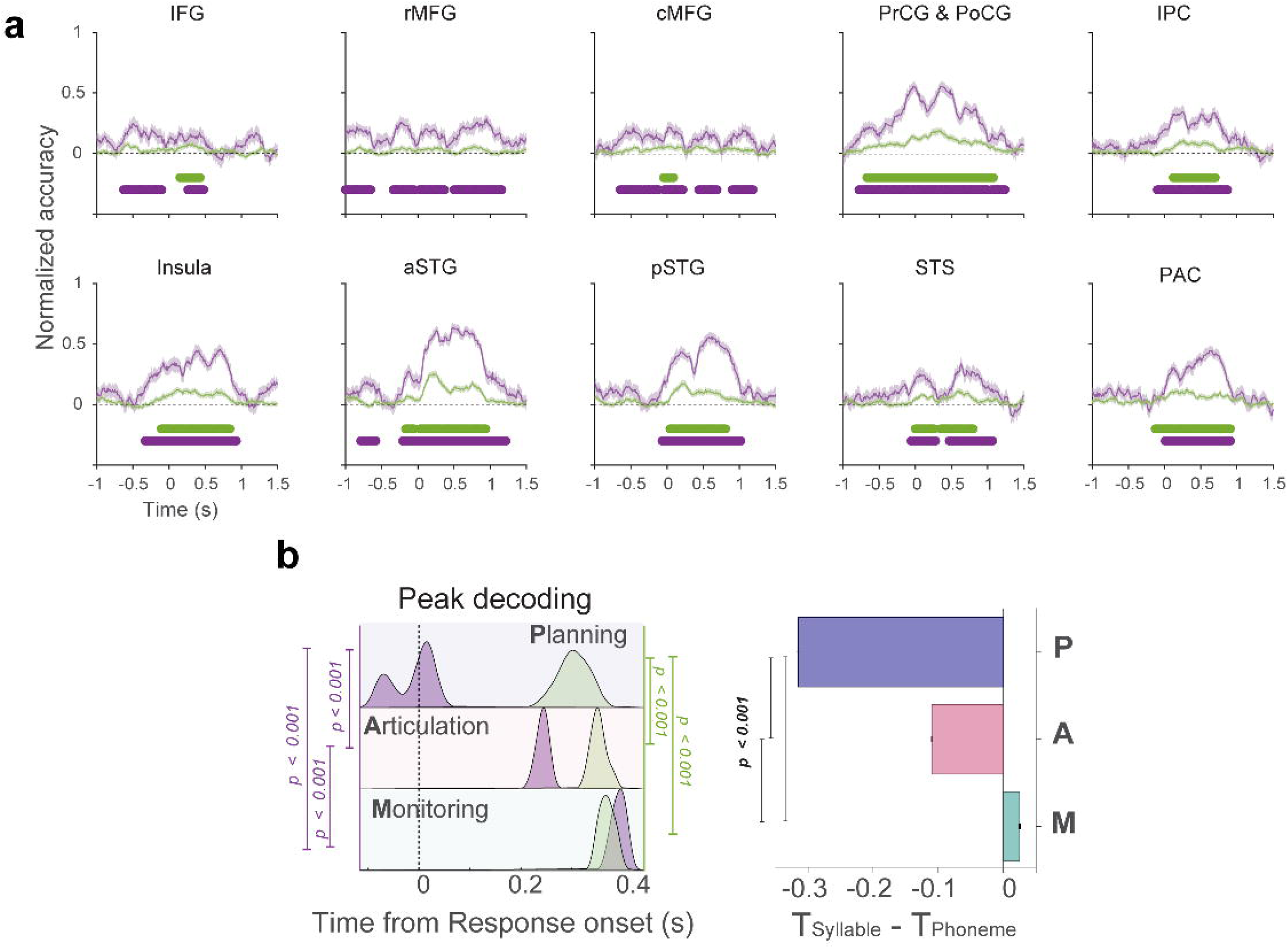

