## Supplementary Figures and Tables for "Distinct neural processes link speech planning and execution"

Duraivel *et al.*

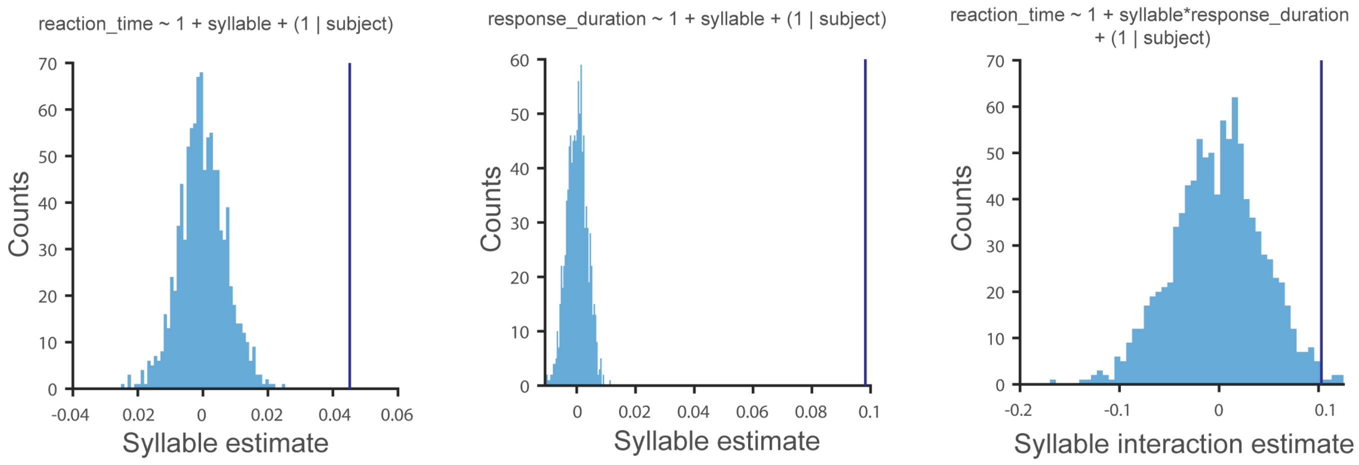


**Supplementary Figure 1**. **Behavior analysis to quantify syllable frame effects on speech planning. Left & Middle**: Linear mixed effects (LME) modeling to quantify the effects of syllable frame on reaction time and duration. Syllable effect (CVC or VCV trials; dark blue) significantly predicts both the reaction time (LME: $t_{(8698)} = 3.7$, β = 0.045 *p* < *0.001*, 95% CI = 0.02 to 0.07, two-sided) and duration (LME: $t_{(8696)} = 10.7$, β = 0.098, *p* < *0.001*, 95% CI = 0.08 to 0.1, two-sided) of the utterance versus a shuffle distribution (n = 1000; light blue histogram). **Right**: LME modeling quantifies the effect of syllable frames on predicting reaction time with response duration as an additional predictor. The interaction term between the syllable frame and the response duration is significant in predicting the reaction time (LME: $t_{(9084)} = 2.75$, β = 0.103, *p* < *0.001*, 95% CI = 0.03 to 0.18, two-sided), indicating that the response duration depends on the identity of syllable frame, highlighting the significant effect of the syllable frame by association. The significant behavior coding provides evidence that speech motor planning reflects the composition of syllable frames.

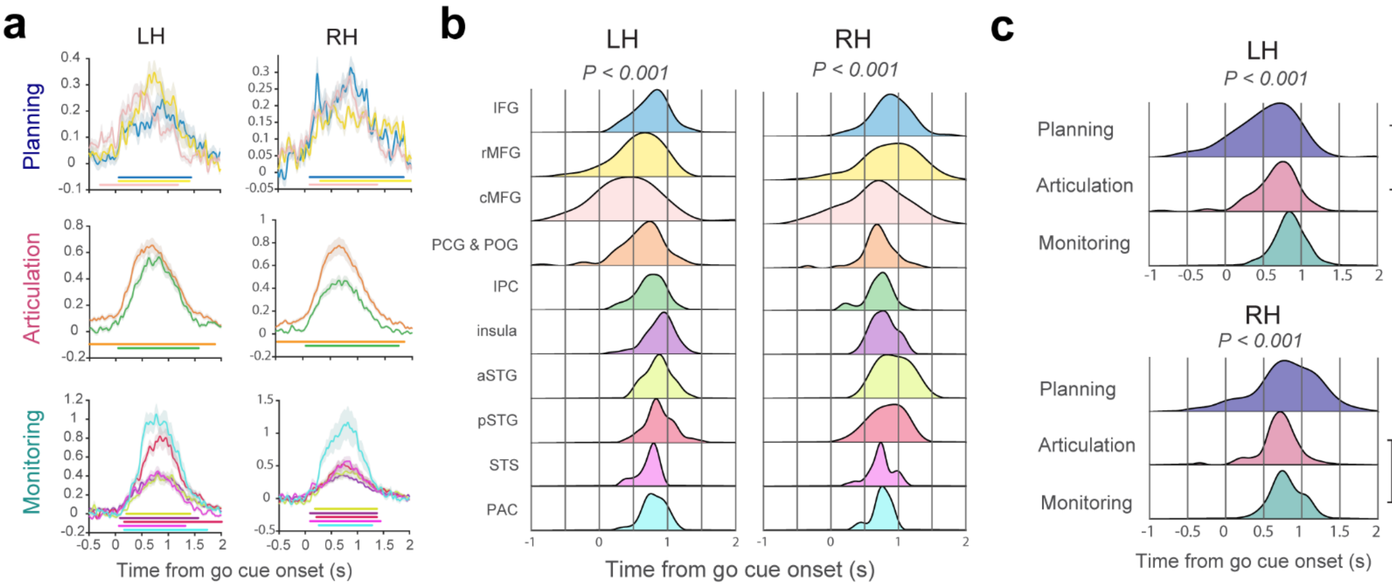


**Supplementary Figure 2**. **Cortical activations from intracranial recordings during speech production aligned to the Go Cue.** **A.** Mean HG activation of ROIs aligned to go cue onset for both left and right hemispheres, averaged within patient. Shaded area denotes standard error. Colored horizontal bars indicate time-periods of significant activation above baseline (one-sided permutation test with cluster correction, cluster forming threshold $p<0.05$, 1000 permutations). **b.** Electrode distributions of peak HG time-points per ROI for each hemisphere with respect to go cue onset. Significant differences in peak activations were observed in both left and right hemispheres, suggesting a temporal progression of neural activations across ROIs around the speech utterance (one-way ANOVAs two-sided: left: $F_{9,717} = 8.03$, $p < 0.001$, right: $F_{9,720} = 6.03$, $p < 0.001$). **c.** Electrode distributions of peak HG time-points per speech network for each hemisphere. This temporal progression was also observed via significant differences in peak activations in both left and right hemispheres (one-way ANOVAs two-sided: left: $F_{2,644} = 52$, $p < 0.001$, right: $F_{2,727} = 8.13$, $p < 0.001$). Post-hoc one-sided t-tests indicate pairwise significance between the speech networks in left hemisphere ($T_{Planning} < T_{Articulation}$, $t_{(376)} = 3.13$, $p = 0.002$; $T_{Articulation} < T_{Monitoring}$, $t_{(504)} = 7.6$, $p < 0.001$, $T_{Planning} < T_{Monitoring}$, $t_{(410)} = 9.7$, $p < 0.001$). Similar significant differences were observed in right hemisphere only for differences between articulation and monitoring networks ($T_{Planning} < T_{Articulation}$, $t_{(529)} = -1.1$, $p = 0.27$ – not significant (NS); $T_{Articulation} < T_{Monitoring}$, $t_{(526)} = 3.3$, $p = 0.001$, $T_{Planning} < T_{Monitoring}$, $t_{(395)} = 1.1$, $p = 0.29$ - NS). Note that the negative t-statistic with planning network denote late neural activations, indicating the monitoring role of right hemispheric pre-frontal regions. These analyses demonstrate the left hemispheric temporal progression of neural responses for planning, articulation, and monitoring networks for speech production. Note that some planning activation is seen prior to the go cue, suggesting a potential involvement of verbal working memory.

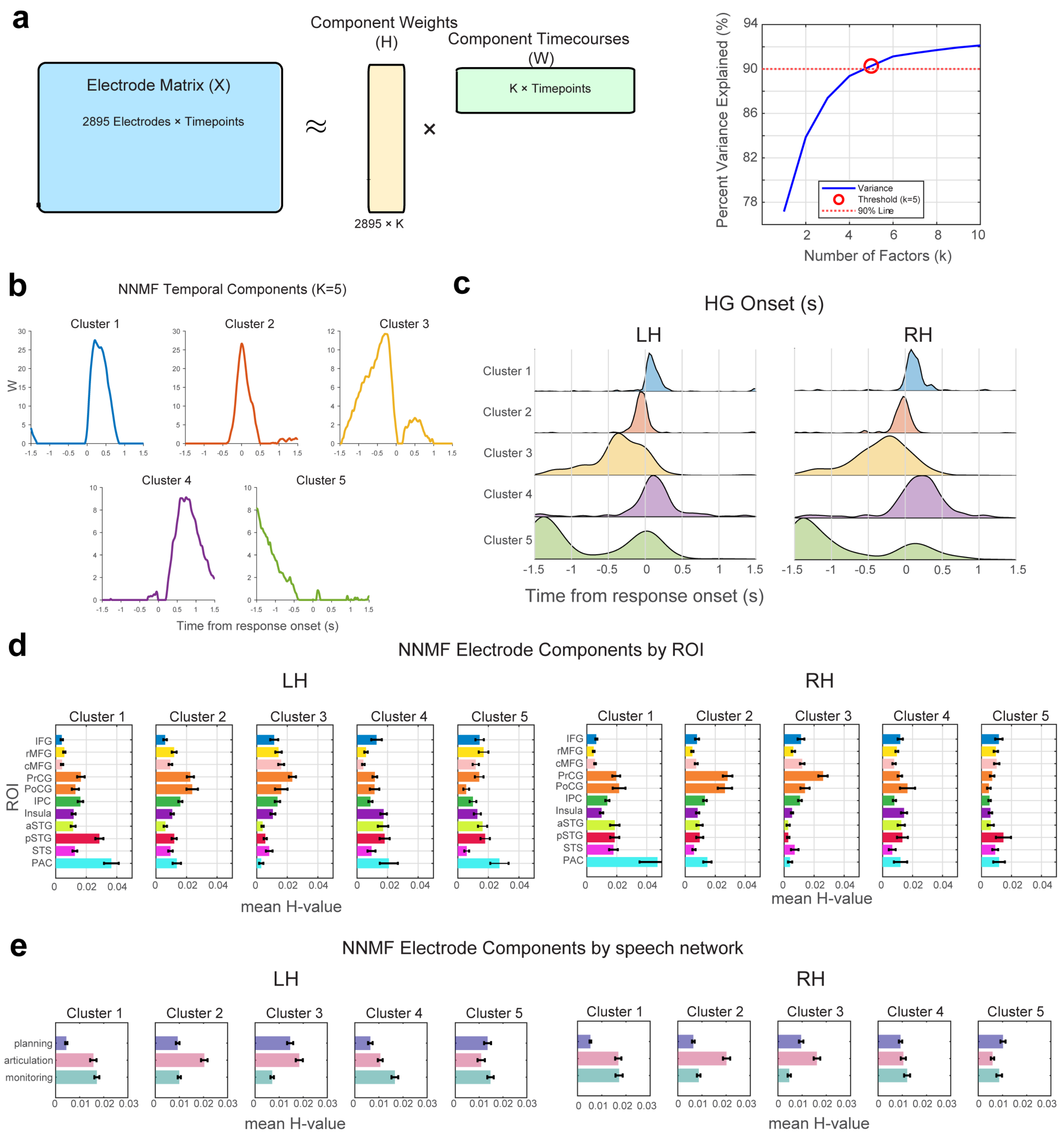


**Supplementary Figure 3. Data driven neural decompositions validate the ROI correspondence of planning and execution process a.** Non-negative matrix factorization schematic to identity the neural sub-components of speech production process. The electrode matrix, containing z-scored HG activations were subjected to matrix factorization, to identify the neural factors (H) that explain the components (W) of neural activations. The resultant variance explained by the model factors were used to determine the number of components. The cumulative variance in percentage (blue) was calculated for different number of components, and the cardinal that explains the top 90% of the variance were chosen to be the optimal candidate (k = 5). **b.** The response time-courses of top 5 clusters that explain components of speech production from planning to execution. The temporal profiles indicate prototypical activation patterns of hypothesized stages of speech production: 1) early auditory monitoring, 2) articulation, 3) planning, 4) late auditory monitoring, 5) verbal working memory. Note – The cardinality of cluster order does not chronologically indicate the stages of speech production. **c.** Electrode distributions of HG onsets per cluster for each hemisphere with respect to response onset. Differences in activation onsets were observed in both left and right hemispheres, suggesting a temporal progression of neural activations around the speech utterance. **d,e.** The electrode factors (H) varied with respect to both ROI and individual speech network, across hemispheres. Error bars indicate mean (± SEM) of factor weights. Cluster 1 and 4 demonstrated maximum activations for monitoring. Cluster 2 signified articulation regions. Finally, left hemispheric cluster 3 demonstrated preference towards the planning activations. Cluster 5 indicate bi-lateral profiles for verbal working memory.

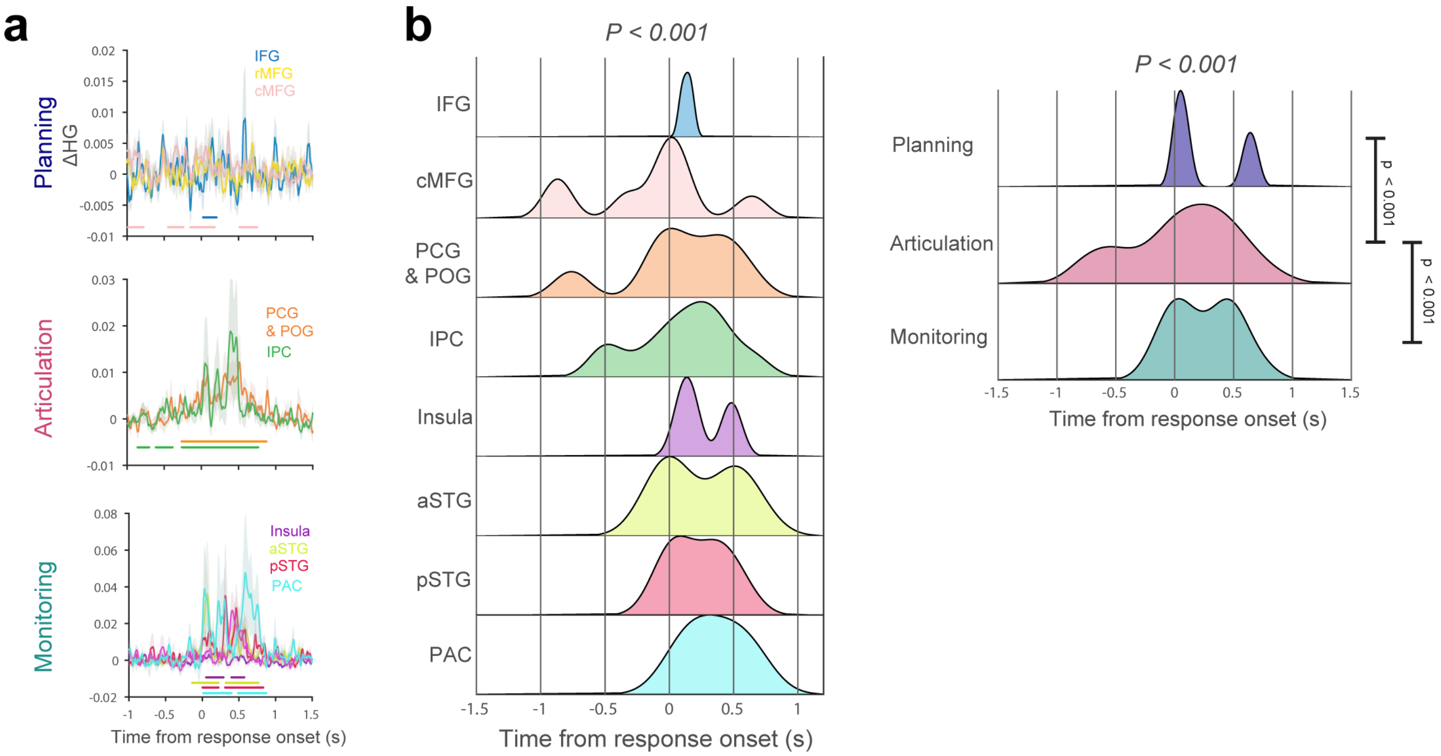


**Supplementary Figure 4**. **Neural code for syllable structures in the right hemisphere. a.** Syllabic frames per ROI show time points with significant syllable discrimination around the utterance. Plots show mean ΔHG contrasts of VCV versus CVC frame for each ROI in the right hemisphere aligned to the response onset. Shaded area denotes standard error. Horizontal bars indicate durations of significant contrasts (across vs. within frame, one-sided permutation test with cluster correction, cluster forming threshold $p<0.05$, 1000 permutations). **b.** Histograms show electrode distributions of peak ΔHG contrasts per ROI & per network. Significant differences in peak ΔHG indicate temporal distinction in syllable activations corresponding to syllable frames, around the utterance (one-way ANOVA two-sided, $F_{7,442} = 6.8$, $p < 0.001$). Significant differences in peak ΔHG for speech networks indicate the temporal distinction of syllable activations across speech networks (one-way ANOVA, two-sided, $F_{2,537} = 19.5$, $p < 0.001$). Post-hoc one-sided t-tests indicate pairwise significance in syllabic frame contrasts (ΔHG) between the networks ($T_{Planning} < T_{Articulation}$, $t_{(406)} = -5.4$, $p < 0.001$; $T_{Articulation} < T_{Monitoring}$, $t_{(415)} = 4.6$, $p < 0.001$) with no significant difference between the planning and monitoring networks, $T_{Planning} < T_{Monitoring}$, $t_{(299)} = 0.16$, $p < 0.001$). Note that the negative t-statistic with planning network denote late processing of syllable frames, possibly indicating the monitoring role of right hemispheric pre-frontal regions. These analyses demonstrate the temporal progression of the neural coding of the syllabic frame within the execution network (articulation to monitoring) in right hemispheric speech ROIs. The hemispheric distinction supports the role of right hemisphere in only executing the planned syllable frames for an utterance.

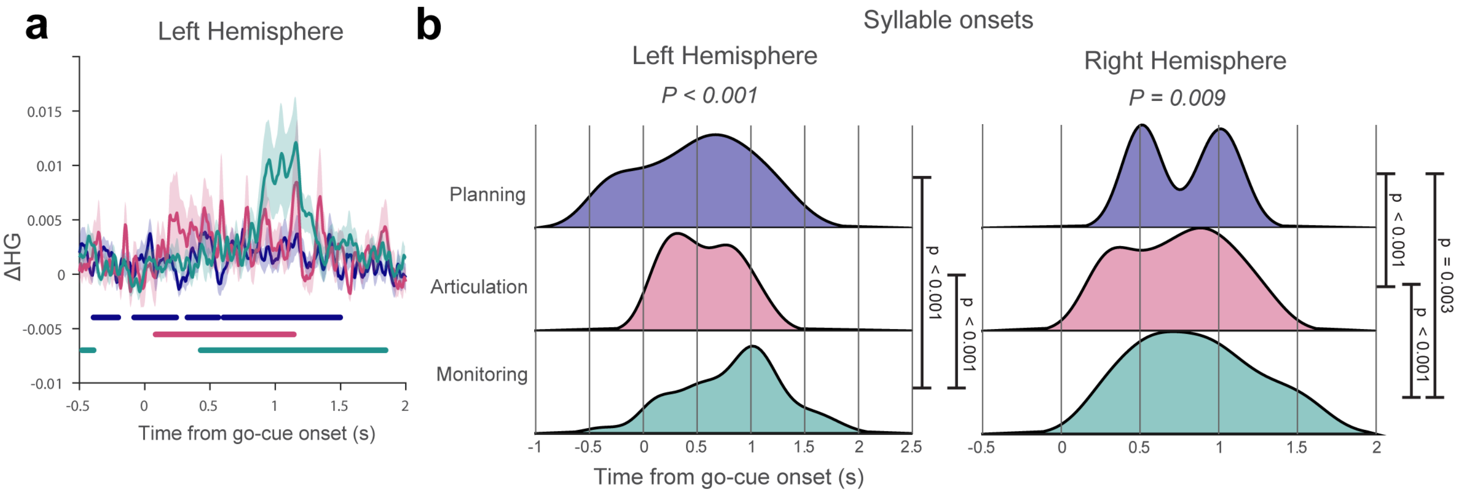


**Supplementary Figure 5**. **Neural code for syllables aligned to go-cue onset. a.** Contrasting syllables for each speech network shows time points with significant syllable discrimination aligned to go-cue onset from the left hemisphere. Plots show mean ΔHG contrasts of VCV versus CVC frame for each speech network. Shaded area denotes standard error. Horizontal bars indicate durations of significant contrasts (across vs. within frame, one-sided permutation test with cluster correction, cluster forming threshold $p<0.05$, 1000 permutations). **b.** Histograms show left hemispheric and right hemispheric electrode distributions of peak ΔHG contrasts for speech network. Significant differences in peak ΔHG show a temporal distinction in syllable activations corresponding to syllable frames, around the go-cue onset (one-way ANOVA two-sided, $F_{2,521} = 26$, $p < 0.001$), with some significant coding during the delay, prior to go-cue onset (a potential indicator of the involvement of verbal working memory). Post-hoc one-sided t-tests show pairwise significance in syllabic frame contrasts (ΔHG) between the networks ($T_{Planning} < T_{Articulation}$, $t_{(314)} = 1.2$, $p = 0.23$ – not significant; $T_{Articulation} < T_{Monitoring}$, $t_{(371)} = 6.3$ $p < 0.001$, $T_{Planning} < T_{Monitoring}$, $t_{(347)} = 6.1$ $p < 0.001$). In right hemisphere, significant differences in peak ΔHG indicate temporal distinction in syllable activations corresponding to syllables, around the utterance (one-way ANOVA two-sided, $F_{2,495} = 4.8$, $p = 0.009$). Post-hoc one-sided t-tests show pairwise significance in syllable contrasts (ΔHG) between the articulation and monitoring networks ($T_{Planning} < T_{Articulation}$, $t_{(339)} = -8.1$, $p < 0.001$; $T_{Articulation} < T_{Monitoring}$, $t_{(378)} = 6.1$, $p < 0.001$, $T_{Planning} < T_{Monitoring}$, $t_{(255)} = -3$, $p = 0.003$), supporting the execution role of right hemisphere. Again, note that the negative t-statistic with planning network denote late processing of syllables, possibly indicating the monitoring role of right hemispheric pre-frontal regions. These analyses demonstrate the temporal progression of the syllabic frame across left hemispheric speech network is preserved with respect to the go-cue, and the left hemispheric planning of syllable frames start prior to instructional go-cue. These results also support the right hemisphere's involvement in execution post go-cue.

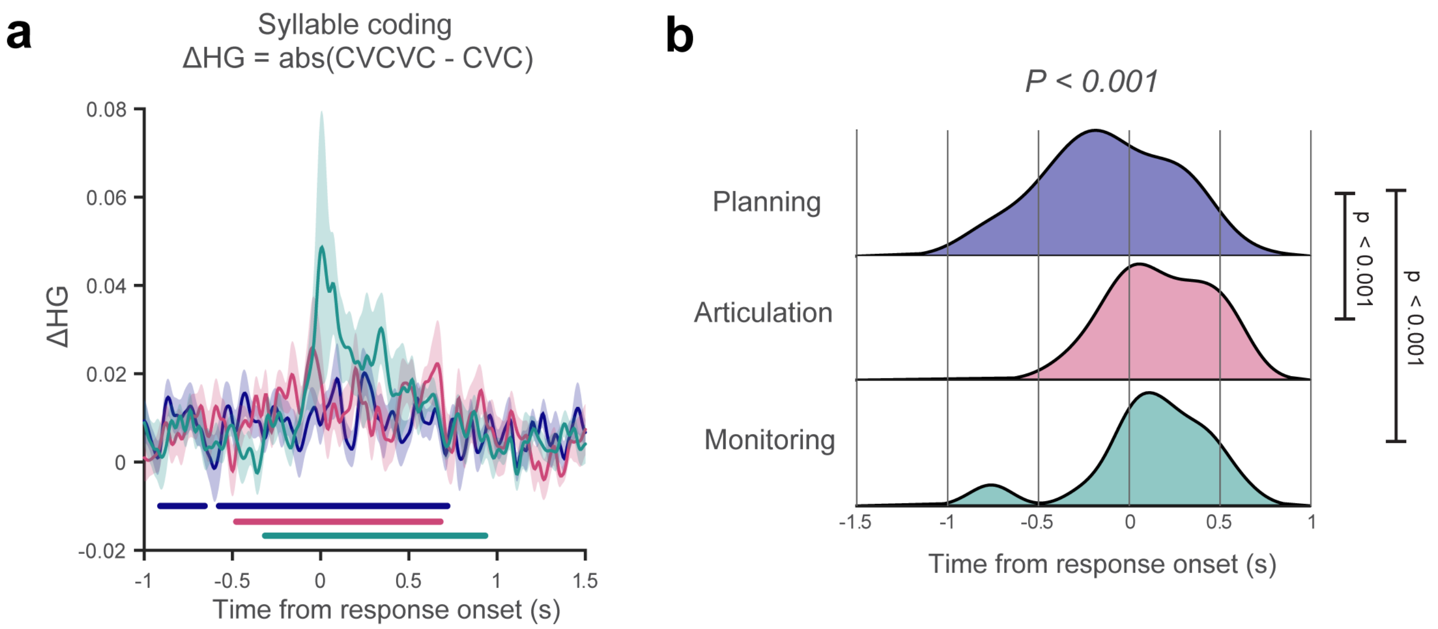


**Supplementary Figure 6**. **Neural code for syllable structure is specific to number of syllables in the structure. a.** Contrasting syllabic frames (CVCVC vs. CVC) per speech network show time points with significant syllable discrimination in the left hemisphere aligned to the utterance. In a subset of patients, a control analysis was performed to rule out the effect of differences in initial phoneme (C vs. V) in the main analysis. Plots show mean ΔHG contrasts of CVCVC versus CVC frame for each speech network aligned to the response onset. Shaded area denotes standard error. Horizontal bars indicate durations of significant contrasts (across vs. within frame, one-sided permutation test with cluster correction, cluster forming threshold *p<0.05*, 1000 permutations). **b.** Histograms show electrode distributions of peak ΔHG contrasts per speech network. Significant differences in peak ΔHG support a temporal progression in syllable activations corresponding to syllable frames, around the utterance (one-way ANOVA two-sided, $F_{2,254} = 18.3$, $p < 0.001$). Post-hoc one-sided t-tests show pairwise significance in syllabic frame contrasts (ΔHG) between the networks ($T_{Planning} < T_{Articulation}$, $t_{(155)} = 3.97$, $p < 0.001$; $T_{Planning} < T_{Monitoring}$, $t_{(182)} = 5.6$, $p < 0.001$), except for the differences between articulation and monitoring network ($t_{(169)} = 0.88$, $p = 0.379$). These analyses indicate the temporal progression of the syllabic frame across left hemispheric speech network is specific to number of syllables in the syllable frame, rather than being driven by phoneme specific information (C vs. V) within the syllable frame.

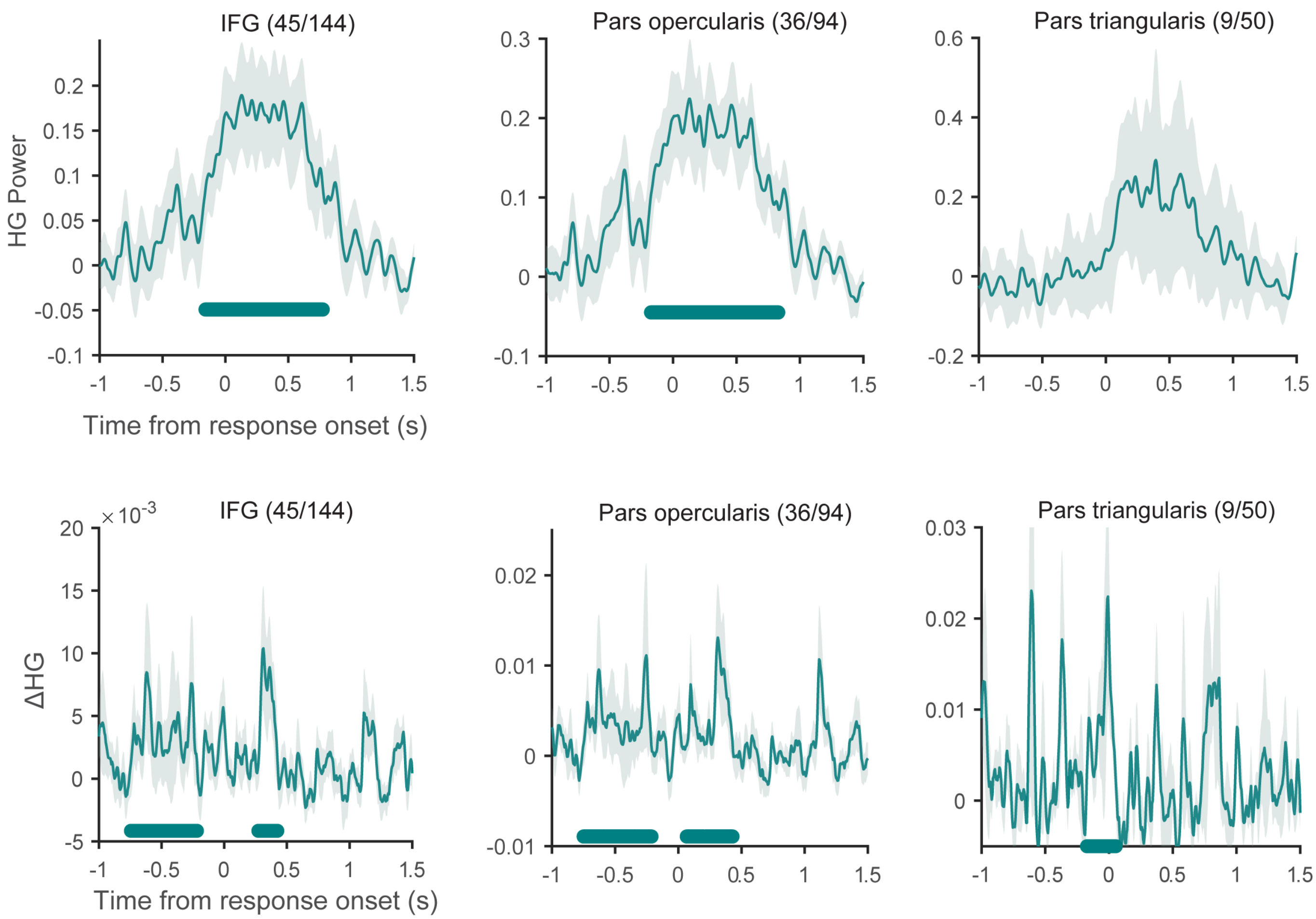


**Supplementary Figure 7**. **Neural activations in left hemispheric IFG are specific to pars opercularis. Top row:** Mean HG activations around response onset for left hemisphere inferior frontal gyrus (IFG), pars opercularis, and pars triangularis. Shaded area denotes standard error. Colored horizontal bars indicate time-periods of significant activation above baseline (one-sided permutation test with cluster correction, cluster forming threshold $p<0.05$, 1000 permutations). Numbers indicate the total number of electrodes with significant HG activations compared to the total number of electrodes within the ROI across patients. **Bottom row:** Contrasting syllabic frames (VCV vs. CVC) per speech network shows time points with significant syllable discrimination around the utterance. Plots show mean ΔHG contrasts of VCV versus CVC frame for each speech network aligned to the response onset. Shaded area denotes standard error. Horizontal bars indicate durations of significant contrasts (across vs. within frame, one-sided permutation test with cluster correction, cluster forming threshold $p<0.05$, 1000 permutations). These results show that neural activations and neural code for syllable frames (ΔHG) from IFG are mainly confined to pars opercularis.

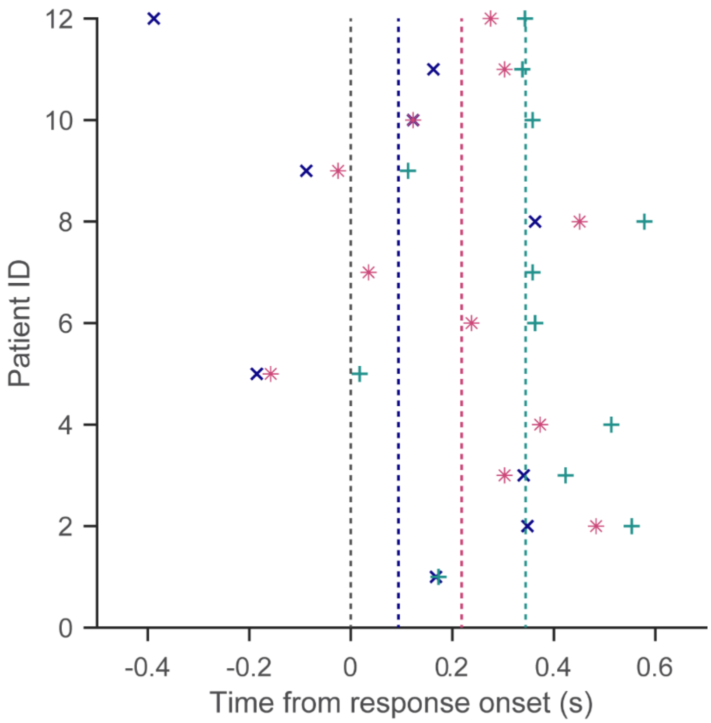


**Supplementary Figure 8**. **The temporal progression of syllable neural code is observed on a patient-specific level.** Scatter plot indicates distribution of peak ΔHG time points of planning (x), articulation (*), and monitoring (+) networks for each patient during utterance. Data is shown for the subset of patients that had significant electrodes in planning along with articulation and/or monitoring regions. Each scatter point denotes the peak ΔHG contrasts of VCV versus CVC frame for each speech network, averaged across electrodes within the patient. The dotted vertical lines indicate average onsets for planning, articulation, and monitoring networks across patients, demonstrating the correct order of temporal progression. The analysis indicates the progression of neural activations from planning to execution of speech is present on a patient specific scale.

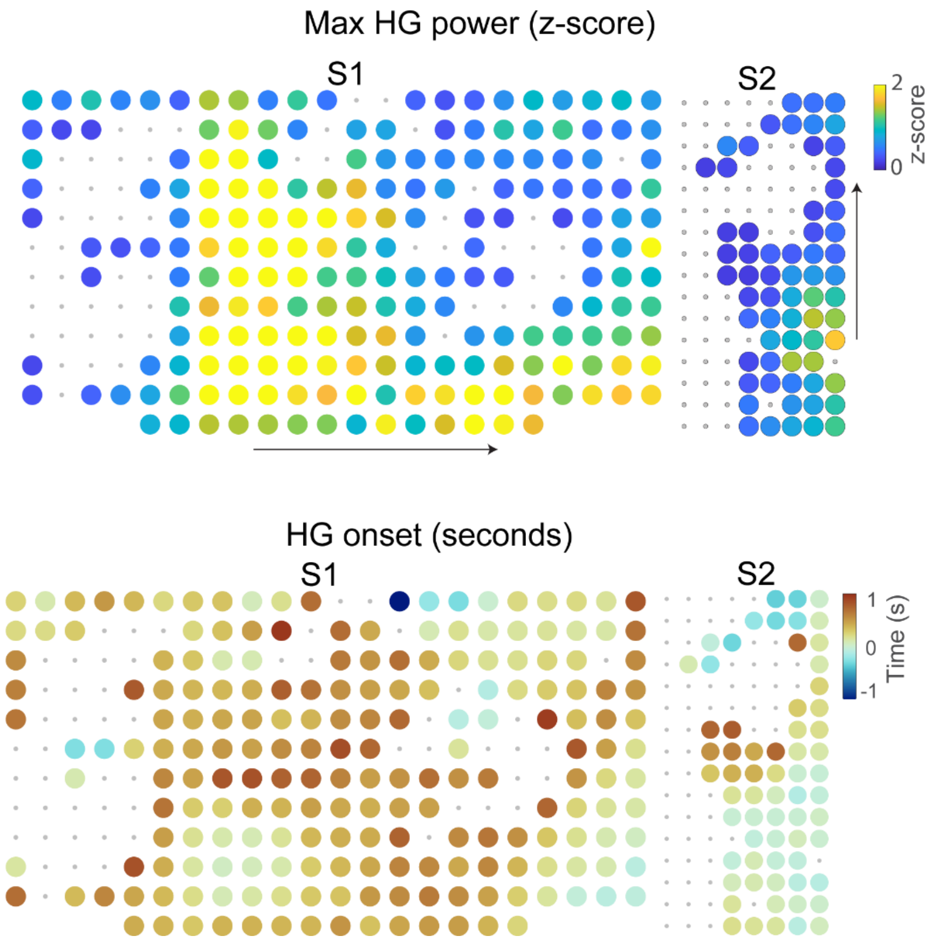


**Supplementary Figure 9**. **µECoG captures spatio-temporally selective activations for speech planning to execution.** High-density µECoG electrode arrays were implanted over planning (anterior) and execution (posterior) networks in two awake patients. Electrode arrays had either 256 electrodes with 1.72 mm inter-electrode distance for S1 (green) or 128 electrodes with 1.33 mm inter-electrode distance for S2 (pink). **Top:** Peak HG activations show fine-scale spatio-temporal patterns during response onset. Large circles indicate electrodes with significant HG activations with heatmap colors corresponding to signal power (FDR-corrected permutation test, $p < 0.05$, 1000 permutations, S1: 201/256 significant/total electrodes, S2: 63/128). Non-significant electrodes are denoted by grey dots. Arrows indicate electrode array orientation from anterior to posterior. **Bottom:** Spatial layouts of HG onsets show spatially varying temporal characteristics for both S1 and S2; 18.9% of significant electrodes reached peak signal power prior to response onset, indicating neural activations for speech planning at the micro-scale electrode level (S1: 12/201, S2: 38/63).

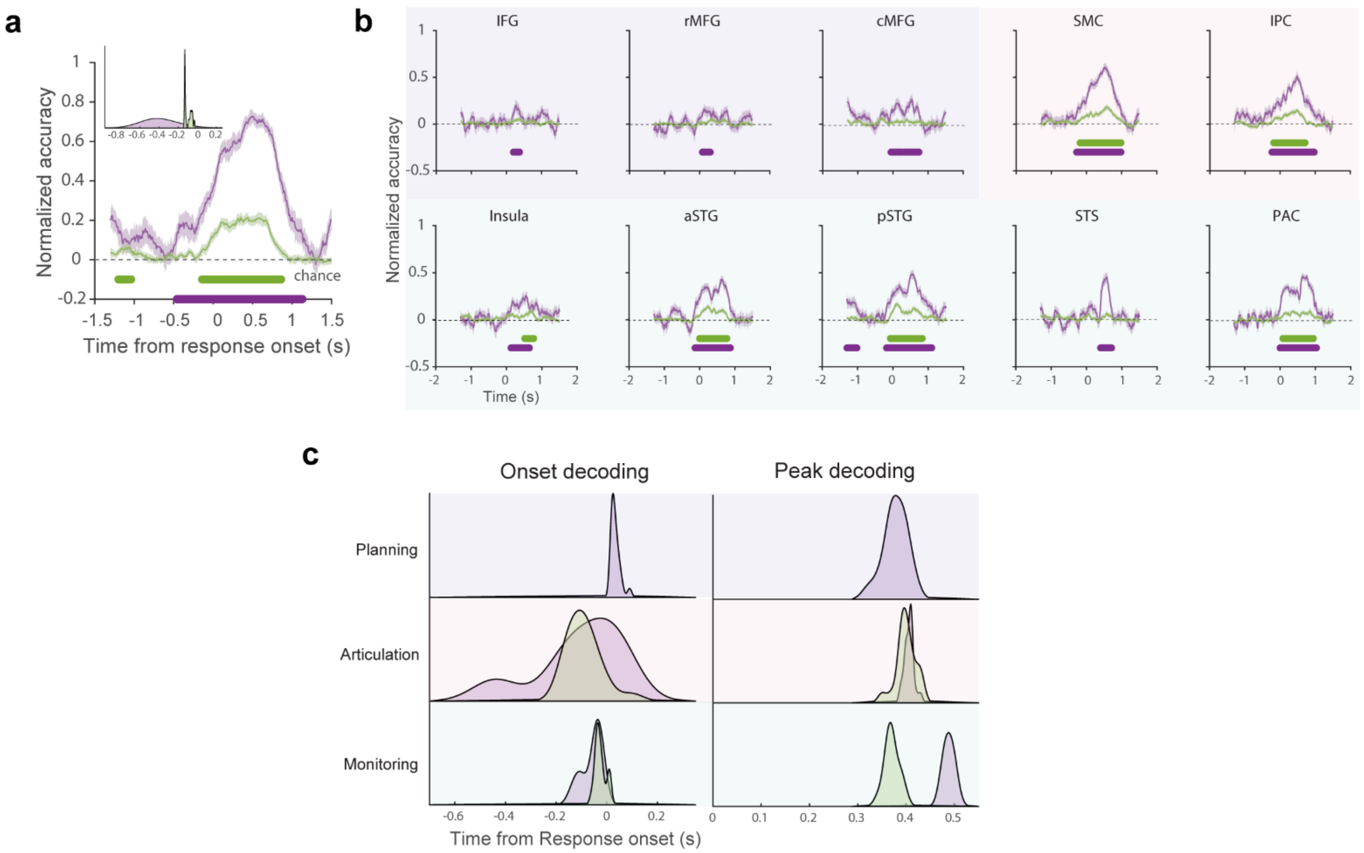


**Supplementary Figure 10**. **Right hemispheric ROIs show significant decoding of syllables and phonemes without a clear hierarchical organization. a.** The temporal evolution of syllables and phoneme activations were obtained using time-resolved neural decoding (200 ms window; 10 ms step size). Solid lines indicate average across 20 instances of decoding iterations, and shaded bars indicate standard errors across instances. Grey traces indicate mean and standard errors of chance decoding which was obtained from randomly shuffling syllable and phoneme labels (250 shuffled decoding models). Horizontal bars indicate durations of significant decoding (one-sided temporal permutation cluster test, cluster forming threshold $p < 0.05$; 250 shuffled decoding models). Colored histograms indicate distributions of decoding onsets of syllables (violet) and phonemes (green), with significant differences between the onsets (one-sided Mann-Whitney U test, $p < 0.001$). **b.** Temporal decoding models also demonstrate a similar syllable and phoneme decoding for each ROI (except phoneme decodes in planning regions and STS). Dotted lines indicate chance decoding. Horizontal bars indicate durations of significant decoding. Background shades indicate ROI networks for planning, articulation and monitoring. **c.** Colored histograms indicate distributions of onset and peak decoding times for both syllable and phoneme for each ROI network. No significant differences in decoding distributions between syllable and phoneme in speech networks. This analysis demonstrates that although the syllables are represented earlier than phonemes on a hemispheric level, the hierarchical organization is absent on a network level.

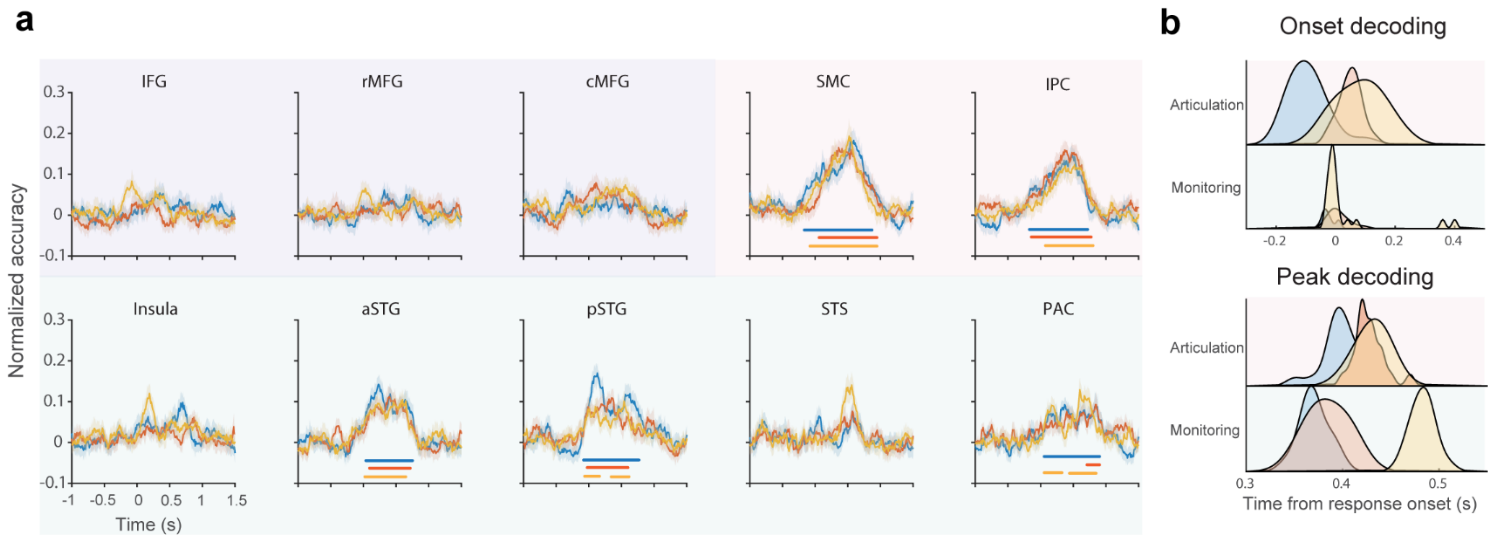


**Supplementary Figure 11**. **Phonological sequence execution in the right hemisphere. a.** Time-resolved decoding models were trained on neural segments aligned to the utterance to decode phonological units at each position within the pseudo-word. Each colored trace indicates temporal evolution of positional phonological units in time for an ROI (mean across 20 decoding instances). Results are shown for all ROIs in the right hemisphere, grouped by planning, articulation, and monitoring networks. Significant phonological unit decoding for all positions was observed only in execution ROIs (articulation and monitoring – except insula and STS). Horizontal bars indicate durations of significant decoding (one-sided temporal permutation cluster test, cluster forming threshold $p < 0.05$; 250 shuffled decoders). Background shading indicates ROI networks for planning, articulation and monitoring. **b.** Colored histograms show distribution times (decoding onsets and peaks) for each positional phonological unit within the ROI network. Positional phonological unit codes were decoded in the correct serial order (except onset decoding for monitoring ROIs, pairwise one-sided Mann-Whitney U test, $p < 0.05$). The presence of ordering demonstrates the serial execution of positional phonological unit codes by the right hemispheric execution network.

**Supplementary Table 1:**

**Clinical summary of patients: Pre-operative epileptic monitoring**

| Patient | Diagnosis | Electrode type: Total No. of recording electrodes | Number of electrodes with significant HG activation |
|---|---|---|---|
| **D20** | Epilepsy | PMT SEEG: 120 | 7 |
| **D22** | Epilepsy | PMT SEEG: 100 | 0 |
| **D23** | Epilepsy | PMT SEEG: 121 | 77 |
| **D24** | Epilepsy | Ad-Tech ECoG w strips: 52 | 12 |
| **D25** | Epilepsy | PMT SEEG: 168 | 6 |
| **D28** | Epilepsy | PMT SEEG: 108 | 44 |
| **D29** | Epilepsy | PMT SEEG: 140 | 60 |
| **D31** | Epilepsy | PMT SEEG: 160 | 51 |
| **D35** | Epilepsy | PMT SEEG: 174 | 52 |
| **D39** | Epilepsy | PMT SEEG: 240 | 107 |
| **D40** | Epilepsy | PMT SEEG: 188 | 4 |
| **D41** | Epilepsy | PMT SEEG: 274 | 118 |
| **D42** | Epilepsy | PMT SEEG: 176 | 27 |
| **D45** | Epilepsy | PMT SEEG: 202 | 54 |
| **D49** | Epilepsy | PMT SEEG: 210 | 113 |
| **D53** | Epilepsy | PMT SEEG: 158 | 11 |
| **D54** | Epilepsy | PMT SEEG: 200 | 0 |
| **D55** | Epilepsy | PMT SEEG: 190 | 64 |
| **D56** | Epilepsy | PMT SEEG: 128 | 54 |
| **D57** | Epilepsy | PMT SEEG: 178 | 85 |
| **D58** | Epilepsy | PMT SEEG: 244 | 67 |
| **D59** | Epilepsy | PMT SEEG: 184 | 63 |
| **D60** | Epilepsy | PMT SEEG: 252 | 111 |
| **D61** | Epilepsy | PMT SEEG: 232 | 9 |
| **D63** | Epilepsy | PMT SEEG: 274 | 77 |
| **D64** | Epilepsy | PMT SEEG: 246 | 36 |
| **D65** | Epilepsy | PMT SEEG: 220 | 101 |
| **D66** | Epilepsy | PMT SEEG: 182 | 44 |
| **D67** | Epilepsy | Ad-Tech ECoG w strips: 72 | 22 |
| **D68** | Epilepsy | PMT SEEG: 146 | 63 |
| **D69** | Epilepsy | Ad-Tech SEEG: 132 | 10 |
| **D70** | Epilepsy | Ad-Tech SEEG: 202 | 0 |
| **D71** | Epilepsy | PMT SEEG: 162 | 49 |
| **D72** | Epilepsy | PMT SEEG: 214 | 173 |
| **D73** | Epilepsy | Dixi SEEG: 200 | 59 |
| **D75** | Epilepsy | Dixi SEEG: 215 | 126 |
| **D76** | Epilepsy | Dixi SEEG: 213 | 145 |
| **D77** | Epilepsy | Dixi SEEG: 157 | 91 |
| **D79** | Epilepsy | Dixi SEEG: 256 | 69 |
| **D81** | Epilepsy | Dixi SEEG: 222 | 111 |
| **D82** | Epilepsy | Dixi SEEG: 263 | 152 |
| **D84** | Epilepsy | PMT SEEG: 230 | 201 |
| **D85** | Epilepsy | Dixi SEEG: 247 | 126 |
| **D86** | Epilepsy | PMT SEEG: 284 | 49 |
| **D88** | Epilepsy | PMT SEEG: 184 | 127 |
| **D91** | Epilepsy | Dixi SEEG: 332 | 123 |
| **D92** | Epilepsy | Dixi SEEG: 218 | 83 |
| **D93** | Epilepsy | Dixi SEEG: 210 | 125 |
| **D95** | Epilepsy | Dixi SEEG: 294 | 88 |
| **D96** | Epilepsy | Dixi SEEG: 234 | 86 |
| **D102** | Epilepsy | Dixi SEEG: 227 | 0 |
| **D103** | Epilepsy | Dixi SEEG: 247 | 2 |

**Clinical summary of patients: Intra-operative awake surgery**

| Patient | Diagnosis | Electrode type/ No. of recording electrodes | Electrode material | Number of electrodes with significant HG activation |
|---|---|---|---|---|
| **S1** | Tumor resection | μECoG: 256 | Platinum-Iridium | 201 |
| **S2** | Parkinson’s | μECoG: 128 | Platinum-Iridium | 63 |
| **S3** | Parkinson’s | μECoG: 128 | Gold | 111 |

**Supplementary Table 2: Stimulus labels for speech repetition task**

| CVC | VCV |
|---|---|
| /bab/ | /abæ/ |
| /bæk/ | /abi/ |
| /bak/ | /æba/ |
| /bup/ | /æbi/ |
| /gab/ | /æbu/ |
| /gæb/ | /æga/ |
| /gæv/ | /æka/ |
| /gak/ | /æpi/ |
| /gav/ | /aka/ |
| /gig/ | /aku/ |
| /gip/ | /ava/ |
| /gub/ | /avæ/ |
| /kab/ | /ibu/ |
| /kæg/ | /ika/ |
| /kub/ | /ikæ/ |
| /kug/ | /ipu/ |
| /pæk/ | /iva/ |
| /pæp/ | /ivu/ |
| /pæv/ | /uba/ |
| /puk/ | /uga/ |
| /pup/ | /ugæ/ |
| /væk/ | /ukæ/ |
| /væg/ | /upi/ |
| /vip/ | /upu/ |
| /vug/ | /uvæ/ |
| /vuk/ | /uvi/ |

**Supplementary Table 3: Anatomical label assignment using Brainnetome labels**

| Network | ROI | Full Anatomical Name | BN Atlas Labels (Left Hemisphere) | BN Atlas Labels (Right Hemisphere) |
|---|---|---|---|---|
| **Planning** | IFG | Inferior Frontal Gyrus | A44d_L, A45c_L, A45r_L, A44op_L, A44v_L | A44d_R, A45c_R, A45r_R, A44op_R, A44v_R |
| **Planning** | rMFG | Rostral Middle Frontal Gyrus | A46_L, A10l_L, A9/46v_L, A9/46d_L | A46_R, A10l_R, A9/46v_R, A9/46d_R |
| **Planning** | cMFG | Caudal Middle Frontal Gyrus | IFJ_L, A8vl_L, A6vl_L | IFJ_R, A8vl_R, A6vl_R |
| **Articulation** | PrCG | Precentral Gyrus | A4hf_L, A6cdl_L, A4ul_L, A4t_L, A4tl_L, A5cvl_L | A4hf_R, A6cdl_R, A4ul_R, A4t_R, A4tl_R, A5cvl_R |
| **Articulation** | PoCG | Postcentral Gyrus | A1/2/3ulhf_L, A1/2/3tonla_L, A2_L, A1/2/3tru_L | A1/2/3ulhf_R, A1/2/3tonla_R, A2_R, A1/2/3tru_R |
| **Articulation** | IPC | Inferior Parietal Cortex | A39c_L, A39rd_L, A40rd_L, A40c_L, A39rv_L, A40rv_L | A39c_R, A39rd_R, A40rd_R, A40c_R, A39rv_R, A40rv_R |
| **Monitoring** | aSTG | Anterior Superior Temporal Gyrus | A38m_L, TE1.0_L, TE1.2_L, A38l_L, A22r_L | A38m_R, TE1.0_R, TE1.2_R, A38l_R, A22r_R |
| **Monitoring** | pSTG | Posterior Superior Temporal Gyrus | A22c_L | A22c_R |
| **Monitoring** | STS | Superior Temporal Sulcus | rpSTS_L, cpSTS_L | rpSTS_R, cpSTS_R |
| **Monitoring** | PAC | Primary Auditory Cortex | A41/42_L | A41/42_R |
| **Monitoring** | Insula | Insula | G_L, vIa_L, dIa_L, vId/vIg_L, dIg_L, dId_L | G_R, vIa_R, dIa_R, vId/vIg_R, dIg_R, dId_R |
